# RudS: Bacterial Desulfidase Responsible for tRNA 4-Thiouridine De-modification

**DOI:** 10.1101/2024.05.15.593862

**Authors:** Rapolas Jamontas, Audrius Laurynėnas, Deimantė Povilaitytė, Rolandas Meškys, Agota Aučynaitė

## Abstract

In this study, we present a comprehensive analysis of a widespread group of bacterial tRNA de-modifying enzymes, dubbed RudS, which consist of a TudS desulfidase fused to a domain of unknown function (DUF1722). RudS enzymes exhibit specific de-modification activity towards the 4-thiouridine modification (s4U) in tRNA molecules, as indicated by our experimental findings. Notably, heterologous overexpression of RudS genes in *Escherichia coli* leads to a significant reduction in tRNA 4-thiouridine content, highlighting the enzyme’s role in tRNA s4U modification regulation. Through a combination of protein modeling, docking studies, and molecular dynamics simulations, we have identified amino acid residues involved in catalysis and tRNA binding. Experimental validation through targeted mutagenesis confirms the TudS domain as the catalytic core of RudS, with the DUF1722 domain facilitating tRNA binding in the anticodon region. Our results suggest a potential role for RudS tRNA modification eraser proteins in prokaryotic tRNA regulation pathways.

**GRAPHICAL ABSTRACT:** 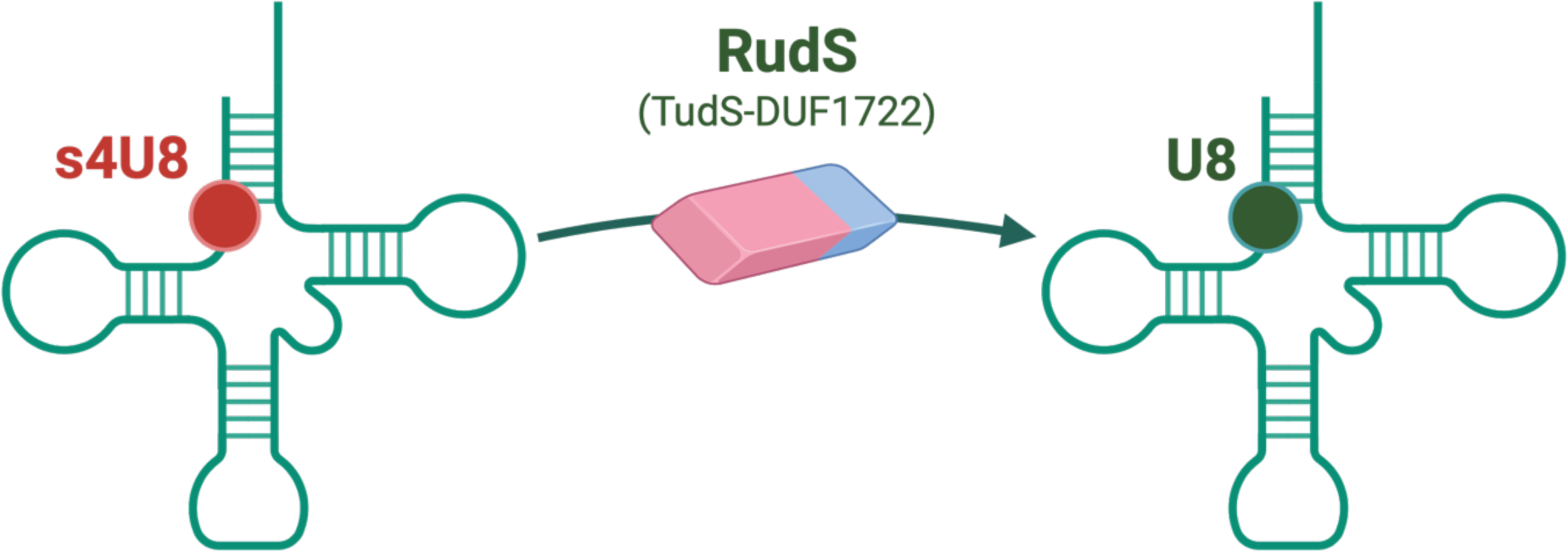

## INTRODUCTION

Sulfur, an element pivotal for life, manifests its biological significance through diverse molecular forms, including the thiol groups of cysteines, thioethers of methionines, various cofactors (e.g., S-adenosyl methionine, [Fe-S] clusters), and as crucial post-transcriptional modifications in transfer RNAs (tRNAs) (1–4). The ubiquity of sulfur-containing tRNA modifications underscores their significance in cellular functions, affecting everything from codon recognition accuracy to thermal stability of tRNA molecules, highlighting their prospective roles in stress responses and cellular regulation (1, 2, 13–19, 5–12).

One of the most abundant sulfur modifications in bacteria and archaea is 4-thiouridine (s4U), most commonly found at position 8 within a tRNA molecule (9, 20). Some tRNA species, however, exhibit a second modification at position 9, resulting in two adjacent s4U modifications (21–23). ThiI, initially identified as an enzyme involved in thiamine biosynthesis in *E. coli*, is typically regarded as the primary biosynthetic enzyme for s4U (24). In fact, enzymes responsible for s4U synthesis vary across species, exhibiting differing structural features and sulfur transfer mechanisms (9, 25–28). Among sulfur-containing tRNA modifications, s4U stands out for its unique susceptibilities, including being a target for UV-A irradiation, which can impair translation by stalling aminoacylation of cross-linked tRNAs (29, 30). Moreover, the s4U modification contributes to thermodynamical stability of tRNA molecules by significantly increasing their melting temperature (22, 31) and affecting the efficiency of incorporation of other modifications (31, 32).

Traditionally viewed as stable entities (33, 34), bacterial tRNAs are now known to undergo rapid degradation under certain conditions, such as amino acid starvation (35), revealing a dynamic balance between tRNA synthesis and degradation (36). This balance is crucial for cellular adaptation to environmental stresses, where tRNA de-modifying enzymes could come into play with a regulatory role, modulating the tRNA pool in response to changing cellular conditions. Several de-modification/eraser enzymes, acting on individual modified nucleosides or nucleobases have been described (37–44), as well as an enzyme which mediates subsequent C-to-U-to-Ψ conversion within tRNA (45). However, up to this date the targeted removal of posttranscriptional modifications from tRNA was only reported for enzymatic demethylation in the context of eukaryotic tRNA homeostasis (46–48).

Despite extensive research into the biosynthesis of thio-modified nucleosides (2, 9, 49–54) the pathways for their enzymatic degradation remain less understood. The dual nature of sulfur, being indispensable yet potentially toxic, necessitates sophisticated cellular mechanisms for its safe storage and transfer. A growing body of evidence suggests that a class of sulfur transferases, including the TudS desulfidase, use a mechanism for direct sulfur transfer via [Fe-S] clusters (49–52, 55–60). We recently provided experimental evidence that TudS enzymes, present in bacteria, archaea, and some eukaryotes, are desulfidases acting on thiolated bases, nucleosides and their monophosphate and triphosphate derivatives with a preference for 4-thio-UMP (41). We also noted that TudS enzymes frequently have additional domains attached to them, potentially leading to a change in substrate preference that allows for the processing of larger molecules, including oligonucleotides or even the tRNA molecule.

Here, we report the discovery of RudS (t<u>R</u>NA 4-thio<u>u</u>ridine <u>d</u>e<u>s</u>ulfidases), a novel class of tRNA de-modifying/eraser enzymes. RudS are fusion proteins combining a TudS desulfidase domain and a domain of unknown function (DUF1722), which specifically target 4-thiouridine (s4U) modification in tRNA, demonstrating a direct role in the cellular management of sulfur-modified tRNAs. This paper delves into the biochemical properties of RudS, elucidating its in vitro and in vivo activities, and sheds light on the molecular interactions critical for its function, thereby contributing to our understanding of the complex regulatory networks governing tRNA stability and function in bacteria.

## MATERIALS AND METHODS

### Bacterial strains, plasmids, and oligonucleotide primers

The bacterial strains used in this study are listed in Supplementary Table 1. The plasmid vectors used in this study are listed in Supplementary Table 2. The oligonucleotide primers used in this study are listed in Supplementary Table 3. Standard techniques were used for DNA manipulations (61). DNA primers were synthesized, and sequencing was performed by Azenta, Germany.

### Construction of bacterial expression vectors

Bacterial vectors for expression in *E. coli* were constructed using the aLICator LIC Cloning and Expression Kit (Thermo Fisher Scientific, USA, #K1291). Bacterial genes were cloned directly from the genomic DNA and the viral gene was synthesized (Thermo Fisher Scientific, USA). Species and genes used in this study are listed in Table 1. Genes were cloned into pLATE11 and pLATE52 expression vectors as described in the manufacturer’s protocol. Genomic DNA was isolated using Quick-DNA Fungal/Bacterial Miniprep Kit (Zymo Research, USA, #D6005), plasmid isolation and PCR product extraction from agarose gels were carried out using GeneJET kits (Thermo Fisher Scientific, USA, #K0503, #K0832). Site directed mutagenesis was carried out using Phusion™ Plus DNA Polymerase (Thermo Fisher Scientific, USA, #F631XL) and a single oligonucleotide primer as described in (62). Codons encoding amino acids of interest were changed to ones either encoding methionine or alanine (63). Empty pLATE11 vector was created by PCR amplification of the backbone and blunt end ligation.

**Table 1:** TudS, RudS and DUF1722 encoding genes used in this study. The conserved cysteine residues are predicted to form the iron-sulfur cluster within the active center of TudS domain.

| Insert | Domain structure | Origin | GeneBank accession ID (82) | Gene locus tag | Conserved cysteines |
| --- | --- | --- | --- | --- | --- |
| RudS_KT | TudS-DUF1722 | *Pseudomonas putida* KT2440 | AE015451 | PP_0741 | Cys17, Cys49, Cys114 |
| RudS_TT | TudS-DUF1722 | *Thermus thermophilus* HB8 | AP008227 | TTHB112 | Cys16, Cys47, Cys111 |
| RudS_ST*<br>(ORF319) | TudS-DUF1722 | *Salmonella enterica* subsp. *enterica* LT2 | AE006468 | STM1389 | Cys12, Cys44, Cys110 |
| RudS_PU | TudS-DUF1722 | *Pseudomonas* sp. MIL9 | JAFEHE010000014 | JQF37_16640 | Cys15, Cys47, Cys112 |
| RudS_PP | TudS-DUF1722 | *Pseudomonas* sp. MIL19 | JAPPVG010000016 | N1078_12140 | Cys18, Cys50, Cys115 |
| RudS_vir | TudS-DUF1722 | *Escherichia* phage 1H12 | NC_049947 | H3V29_gp66 | Cys12, Cys44, Cys110 |
| TudS | TudS | Uncultured bacterium | 6Z92_A (40, 41, 60) | - | Cys10, Cys43, Cys104 |
| YbgA | DUF1722 | *Escherichia coli* DH5α | NZ_SGJA000000000 | b0707 | - |

\*The RudS protein from *Salmonella enterica* (RudS\_ST here) was previously referred to as ORF319 in the literature (83).

### Cultivation of bacteria and overexpression of recombinant proteins

For the bacterial growth assay, *E. coli* HMS174 Δ*pyrF* strains, either carrying the pLATE11 vector with an insert or an empty pLATE11 vector as a negative control, were initially cultured in LB medium supplemented with 100 µg/mL ampicillin overnight at 37 °C, 180 rpm. The overnight cultures were subsequently diluted in M9 minimal media (1× M9 salts, 1 mM MgSO_4_, 0.05 mM CaCl_2_, 0.4% glucose, 100 µg/mL ampicillin, 0.1 mM IPTG) uniformly to OD_600_ 0.001. If necessary, inoculates were supplemented with nucleobases at the final concentration of 200 µM, and 150 µL of suspensions were dispensed into the wells of 96-well flat-bottom plate. Bacterial growth was monitored using Infinite M200 PRO (Tecan, Switzerland) microplate reader. Plates were incubated at 37 °C with periodic shaking for 30 s every 5 min and OD_600_ measurements every 15 min.

For tRNA isolation, a single colony of *E. coli* BL21(DE3), either carrying the pLATE11 vector with an insert or an empty pLATE11 vector as a negative control, was inoculated into 20 mL of LB medium supplemented with 100 µg/mL ampicillin. Cultures were incubated at 37°C, 180 rpm. Upon reaching the log phase (OD_600_ ∼0.6), production of recombinant protein was induced by the addition of IPTG to a final concentration of 0.1 mM. Cells were harvested by centrifugation 4 h post-induction and used for tRNA isolation immediately or stored at -20°C until further use.

For protein purification, an auto-induction medium with slight modifications was used (64). The base of the medium consisted of 10 g tryptone, 10 g yeast extract, 10 g glycerol per 1 L of medium. Medium was supplemented with 50× salt solution (1.25 M Na_2_HPO_4_, 1.25 M KH_2_PO_4_, 2.5 M NH_4_Cl, 0.25M Na_2_SO_4_), 50× lactose solution (25% glycerol, 2.5% glucose, 10% α-lactose), MgSO_4_ (2 mM final concentration), 1000× trace metal solution, and 100 µg/mL ampicillin. A single colony of *E. coli* BL21(DE3) carrying pLATE52 vector with an insert was inoculated into 50 mL of medium and incubated at 37 °C, 180 rpm. After 6 h, the temperature was reduced to 20°C, and the incubation continued for additional 16-20 h. To verify induction, 1 µL of a 2% X-Gal solution was added to 1 mL of the culture, followed by incubation for 30 min at 37 °C.

### Preparation of bulk tRNA

Bulk tRNA was prepared with slight modifications to the method described in (65). Harvested *E. coli* BL21(DE3) bacterial culture was resuspended in 600 µL of extraction buffer (1 mM Tris-HCl, pH 7.4, 10 mM Mg-Acetate) and mixed with 600 µL of ROTI Aqua-Phenol (Carl Roth, Germany, #A980.3). The mixture was mixed using vortex mixer at 2000 rpm for 15 min, followed by centrifugation at 30000 *g*, 16°C for 10 min. The aqueous phase was mixed with 0.1 volume of 5 M NaCl and 2 volumes of 100% ethanol, followed by centrifugation at 30000 *g*, 4°C for 10 min. Pellet was resuspended in 550 µL of 1 M NaCl and centrifuged at 30000 *g*, 4°C for 10 min. Subsequently, 500 µL of the supernatant was mixed with 1250 µL of 100% ethanol and incubated at –20°C overnight. The precipitate was collected by centrifugation at 30000 *g*, 4°C for 10 min and dissolved in 25 mM K-phosphate buffer (pH 6.5).

For the purification step, the sample was applied to a HiTrap DEAE Sepharose FF column (Cytiva, USA, #17505501) pre-equilibrated with buffer A (25 mM Na-phosphate pH 6.5, 50 mM NaCl, 5 mM MgSO_4_) using AKTA Pure FPLC system (Cytiva, USA). The tRNA was eluted by applying a linear gradient (0-100%) of buffer B (25 mM Na-phosphate pH 6.5, 1 M NaCl) over 10 min using 10 CV. The eluted fractions were precipitated by the addition of 3 volumes of 100% EtOH. The resulting pellet was resuspended in DEPC-treated water.

### tRNA digestion and analysis of modified nucleosides

For the analysis of modified nucleosides using HPCL-MS/MS, 1 µg of tRNA was heat denatured for 5 min at 95°C and subjected to digestion at 37°C for 16 h using Nucleoside Digestion Mix (New England BioLabs, USA, #M0649S). After digestion, proteins were precipitated by adding an equal volume of acetonitrile to the digested tRNA. The mixture was mixed at 1400 rpm, 37°C for 10 min, and centrifuged at 30000 *g*, 4°C for 20 min. Supernatant was used for nucleoside analysis.

4 μL of the supernatant were analyzed using liquid chromatography-tandem mass spectrometry with a Nexera X2 UHPLC system coupled with LCMS-8050 mass spectrometer (Shimadzu, Japan) equipped with an ESI source. The chromatographic separation was carried out using a 3 × 150 mm YMC-Triart C18 (particle size 3 μm) column (YMC, Japan, #TA12S03-1503WT) at 40°C and a mobile phase that consisted of 0.1% formic acid (solvent A) and acetonitrile (solvent B) delivered in gradient elution mode at a flow rate of 0.45 mL/min. The following elution program was used: 0 to 1 min, 5% solvent B; 1 to 5 min, 95% solvent B; 5 to 7 min, 95% solvent B; 7 to 8 min, 5% solvent B; 8 to 12 min, 5% solvent B. Modified nucleosides were detected using transitions m/z 247→115 (dihydrouridine) and 261→129 (4-thiouridine) at interface temperature of 300°C and desolvation line temperature of 250°. N_2_ was used as nebulizing (3L/min) and drying (10L/min) gas, dry air was used as heating (10L/min) gas. The data were analyzed using LabSolutions LCMS software. For quantification, the amount of 4-thiouridine was normalized to the total dihydrouridine content (66).

### Western blot analysis

In total, 2 mL of induced bacterial cultures (see “Cultivation of bacteria and overexpression of recombinant proteins”) were centrifuged, and the pellets resuspended in 2 mL Tris-HCl pH 7, 250 mM NaCl buffer. Cells were disrupted using ultrasonic disintegrator and centrifuged at 30000 *g*, 4°C for 10 min. The concentrations of the soluble fractions of the crude extracts were measured using Pierce Bradford Assay Kit (Thermo Fisher Scientific, USA, #23246). All samples were adjusted to a concentration of 200 ng/µL, and 2 µg of soluble crude extract was loaded onto a 14% SDS-PAGE gel using the Bio-Rad Mini-PROTEAN electrophoresis system and a 10-well or 15-well, 0.75 mm comb (Bio-Rad, USA, #1653355). Following the electrophoresis, proteins were transferred onto a 0.45 µm nitrocellulose membrane (Thermo Scientific, USA, #88018) and blocked using TBST (10 mM Tris-HCl pH 7.5, 150 mM NaCl, 0.1% Tween 20) containing 0.2% I-Block reagent (Thermo Fisher Scientific, USA, #T2015) for 1 h at room temperature. The membrane was incubated overnight at 4°C with anti-His-Tag antibodies (Thermo Fisher Scientific, USA, #MA1-21315) diluted 1:1000 in the blocking solution. The following day, membranes were incubated with the HRP-conjugated anti-mouse secondary antibody (Carl Roth, Germany, #4759) diluted 1:10000 in blocking solution for 1 h at room temperature. Detection of protein-antibody complexes was performed using an enhanced chemiluminescence substrate (Thermo Fisher Scientific, USA, #32209) and digitally imaged with Azure 280 chemiluminescence detection system (Azure Biosystems, USA). Band intensities were quantified using ImageJ software (National Institutes of Health, USA).

### Purification of RudS_KT recombinant protein

Cells from 50 mL of induced *E. coli* BL21(DE3) cultures (see “Cultivation of bacteria and overexpression of recombinant proteins”) were resuspended in 7.5 mL of buffer A (50 mM TRIS-HCl pH 8, 500 mM NaCl, 10 mM imidazole, 10% (w/v) glycerol), supplemented with ∼0.1 mg of DNase I (Roche, Switzerland, #10104159001), 15 mM MgSO_4_ and 1mM PMSF. Cells were disrupted using an ultrasonic disintegrator and centrifuged at 30000 *g*, 4°C for 10 min. The supernatant was applied to 1 mL HiTrap Chelating HP chromatography column (Cytiva, USA, #17040801) pre-equilibrated with buffer A using AKTA Pure FPLC system (Cytiva, USA).

The protein was eluted by applying linear gradient (0–100%) of buffer B (50 mM TRIS-HCl pH 8, 500 mM NaCl, 500 mM imidazole, 10% (w/v) glycerol) over 10 min using 10 CV. Following elution, the buffer in the fractions was exchanged with buffer S (50 mM TRIS-HCl pH 8, 500 mM NaCl, 10% (w/v) glycerol) using 5 mL HiTrap Sephadex G-25 Desalting columns (Cityva, USA, # 17140801).

To minimize the protein’s exposure to oxygen, all buffers were degassed under vacuum for 30 min with stirring. The fractions were collected in 0.5 mL aliquots in 0.5 mL tubes and sealed immediately after fractionation.

Purified protein samples were analyzed using 14% SDS-PAGE gel, protein concentration was measured using Pierce™ Bradford Protein Assay Kit (Thermo Fisher Scientific, USA, #23200), protein spectrum analysis was carried out using GENESYS™ 150 UV-Vis Spectrophotometer (Thermo Fisher Scientific, USA), iron content in protein samples was determined using ferene (Merck, Germany, #P4272) and FeCl_2_ as a standard (67).

### Determination of RudS_KT molar mass

The molar mass RudS_KT was determined by analytical gel filtration using a Superose 12 10/300 GL column (Cytiva) previously equilibrated with 50 mM TRIS-HCl pH 8, 500 mM NaCl, 10% (w/v) glycerol, under aerobic conditions. Carbonic anhydrase (29 kDa), bovine serum albumin (66 kDa), alcohol dehydrogenase (150 kDa), and β-Amylase (200 kDa) were used as standards (Sigma-Aldrich, USA, #1002033699) for column calibration. The experiment was repeated three times to determine the standard deviation.

### In vitro activity assay

In vitro tRNA 4-thiouridine desulfidation assay was carried out aerobically using purified RudS_KT and total tRNA from *E. coli* MRE 600 (Roche, Switzerland, #10109541001). Purified RudS_KT concentrations were adjusted to 0.7 mg/mL (17.85 µM) with buffer S (50 mM Tris-HCl pH 8, 500 mM NaCl, 10% (w/v) glycerol). Reactions were carried out in 100 µL reaction mixtures containing 5.3 µM RudS_KT, 0.4 µM tRNA, 100 mM Tris-HCl pH 7 and 150 mM NaCl at 22°C. Reactions were stopped by heating the samples at 95°C for 5 min following by centrifugation at 30000 *g*, 4°C for 10 min and ethanol precipitation of tRNA in supernatant. Samples were digested to single nucleosides and analyzed by HPLC-MS/MS as described above.

To test the (p)ppGpp effect on RudS_KT activity, 10 fold molar excess of guanosine-3’,5’-pentaphosphate (pppGpp) (Jena Bioscience, Germany, #NU-885S) or guanosine-3’,5’-tetraphosphate (ppGpp) (Jena Bioscience, Germany, #NU-884S) was added to equimolar amount of tRNA and RudS_KT. Reactions were carried out in 100 µL reaction mixtures at 22°C containing 4 µM RudS_KT, 4 µM tRNA, 100 mM Tris-HCl, pH 7, 150 mM NaCl and 40 µM of (p)ppGpp. Reactions were stopped by heating and analyzed as described above.

### Electrophoretic mobility shift assay (EMSA)

Non-radioactive EMSA was adapted from (68) with slight modifications. Binding reaction was carried out in 20 µL volume and contained 20 mM Tris-HCl, 10% glycerol, 200 mM NaCl, 0.5 µg (1 µM final concentration) total *E. coli* tRNA and 6 µM his-tagged *E. coli* pseudouridine synthase TruB (positive shift-control) or 1–10 µM RudS_KT. After 1 min incubation on ice, whole binding mixture was loaded into 2% agarose gel prepared by using RNase-free TBE buffer (Invitrogen, USA) and DEPC treated water. Gel and TBE running buffer was supplemented with SYBR™ Green II RNA Gel Stain (Invitrogen, USA, S7564). The gel was run at 7 V/cm for 90 min and imaged using Azure 280 fluorescence detection system (Azure Biosystems, USA).

### Modelling and molecular dynamic simulations

The RudS_KT structures were modeled using Alphafold2 (69) and trRosetta (70) web servers. Both methods utilized multiple sequence alignments generated with mmseqs2 (71) and Hhblits (72) for Alphafold2 and trRosetta, respectively. The RudS_KT holoenzyme model was constructed by selecting the best model from Alphafold2 and incorporating the iron-sulfur cluster from PDBID:6z96 (60). Molecular dynamics calculations were performed using the AMBER20 software package (73). The holoenzyme model was parameterized with the AMBER ff14SB force field (74) and the TIP3P explicit solvent model with a 12 Å protein water solvation box and approximately 150 mM NaCl. The angle and bond parameters for the iron-sulfur cluster were adopted from (75), with charges adjusted for different iron oxidation states.

A crystal structure of *E. coli* phenylalanine tRNA (PDBID:6y3g) (76) served as the model substrate, featuring a 4-thiouridine moiety at the 8th position. As the original structure rendered the thiouridine moiety inaccessible to the enzyme, we manually repositioned the base to obtain an initial substrate structure suitable for enzymatic interaction. Additionally, calcium ions in the crystal structure were replaced with magnesium ions. Both the original and repositioned structures were parameterized using identical water models and ranges as the protein, supplemented with OL3 and modrna8 force field for modified RNA (77–79).

The initial enzyme-substrate structures were docked using default settings on the HDOCK web server (80). The resulting structures were ranked based on the distance between the sulfur atom in the thiouridine moiety and the relevant iron atom in the iron-sulfur cluster. The structure that performed best in this regard was selected for further calculations. Molecular dynamics simulations, with various restraints detailed in the Supporting Information, were carried out for the holoenzyme, tRNA substrates, and enzyme-substrate complexes.

## RESULTS

### TudS and RudS proteins do not share the same phenotype in vivo

Our recent research revealed that a family of widespread bacterial proteins consisting of a stand-alone TudS domain (formerly DUF523) salvage thiolated uracil derivatives with the highest preference for thiouridine monophosphate (4-thio-UMP), which most likely emanates from cellular tRNA degradation (41). Although the single-domain TudS proteins are predominant in this family of proteins (9436, or 58.9% of all reported sequences), a significant portion is found fused with DUF1722 domain (6425, or 40.1% of all reported sequences) (81). Notably, the *P. putida* KT2440 genome encodes for both a stand-alone TudS (GenBank gene locus ID: PP_5158) and a TudS-DUF1722 fusion protein RudS (GenBank gene locus ID: PP_0741). Gene knockout studies in *P. putida* KT2440 suggested that contrary to stand-alone TudS (PP_5158) the RudS (PP_0741) gene product does not participate in exogenous 4-thiouracil/4-thiouridine utilization (41), which prompted the question whether a TudS-DUF1722 fusion protein could have an alternative substrate to that of a stand-alone TudS.

To confirm that RudS fusion proteins’ activity in vivo is different from stand-alone TudS, we firstly cloned and overexpressed the *P. putida* PP_0741 gene (RudS_KT) product in uracil auxotrophic *E. coli* HMS174 Δ*pyrF*, replicating conditions of our previous studies (40, 41, 60). Using liquid M9 media we observed the growth restoration on 4-thiouracil (Figure 1D), albeit after an extended lag phase compared to TudS (Figure 1B). No growth was observed on 2-thiouracil, unlike with previously characterized TudS proteins. These findings suggested that RudS_KT exhibits residual activity towards thiolated uracil compounds as substrates.

**Figure 1:**
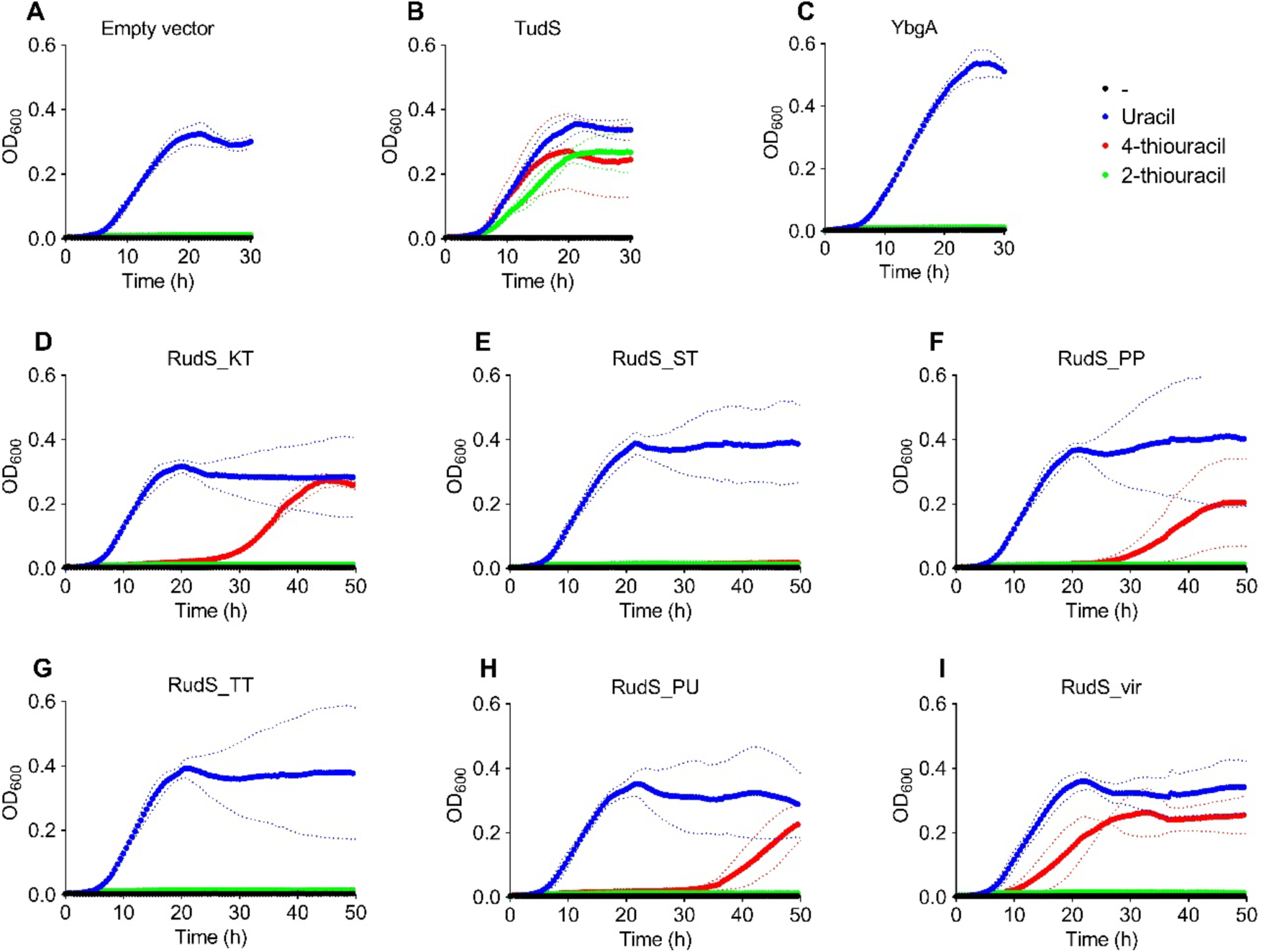
Growth curves of uracil auxotrophic *E. coli* HMS174 Δ*pyrF* expressing TudS, RudS and DUF1722 domain encoding genes in the presence of uracil (blue), 2-thiouracil (green) and 4-thiouracil (red). Dotted lines represent standard deviation. A: Empty vector carrying bacteria (negative control). B: TudS producing bacteria (positive control). C: YbgA (stand-alone DUF1722 domain) producing bacteria. D: RudS_KT producing bacteria. E: RudS_ST producing bacteria. F: RudS_PP producing bacteria. G: RudS_TT producing bacteria. H: RudS_PU producing bacteria. I: RudS_vir producing bacteria.

We cloned four additional RudS genes from common laboratory strains and soil bacteria (Table 1). Notably, *Escherichia coli* lacks the TudS or RudS homologues, but possesses a stand-alone DUF1722 domain encofding gene *ybgA*, which was tested as well (Figure 1C). Ten gene sequences of viral origin RudS were reported, all within the *Siphoviridae* family (81), of which we chose the RudS_vir from *Escherichia* phage 1H12 for further analysis (Figure 1I).

The stand-alone TudS gene product (Figure 1B) served as a positive control for this experiment, while the stand-alone DUF1722 gene product (Figure 1C) exhibited similar behavior to the negative control (Figure 1A). None of the tested RudS variants supported the growth of the uracil-auxotroph in the same manner as TudS. Similarly to RudS_KT (Figure 1D), bacteria with RudS from other *Pseudomonas* species (Figure 1F, H) displayed a significant lag phase before initiating growth on 4-thiouracil, while viral-origin RudS showed the shortest lag phase on 4-thiouracil (Figure 1I). In contrast, the RudS_TT from *Thermus thermophilus* (Figure 1G) and RudS_ST from *Salmonella enterica* (Figure 1E) were unable to rescue the uracil auxotrophic strain on either on 4-or 2-thiouracil. None of the tested RudSes supported the growth on 2-thiouracil. The growth curves suggested that thiolated uracils are not the primary substrates for RudS variants, as they did not exhibit significant activity towards these compounds.

### Heterologous overexpression of RudS genes results in decreassed 4-thiouridine content of *E. coli* tRNA

Observation of either promiscuous or absent TudS-like activity in RudS suggested that TudS and RudS likely have distinct functions in vivo. However, the presence of the TudS domain within RudS implies a potential role related to 4-thiouridine desulfidation. This assumption is supported by the fact that genes encoding the DUF1722 domains, are conserved in prokaryotes, as well as are the genes encoding ThiI, the enzyme responsible for s4U synthesis. Additionally, the phylogenetic analysis using PhyloCorrelate (84) of bacterial and archaeal genes revealed a notable pattern: most species with the DUF1722 domain gene also encode ThiI gene (94% of DUF1722 encoding species), although not vice versa (Supplementary Figure S1) indicating a putative link between these genes. Given that 4-thiouridine, to the best of our knowledge, is conserved in tRNA, we decided to overexpress the RudS homologs in *E. coli*, isolate bulk tRNA, and examine its 4-thiouridine content.

Genes listed in Table 1 were overexpressed in *E. coli* BL21(DE3) cells, total tRNA was extracted from these cells, and the level of s4U in it was quantified using HPLC-MS/MS. A substantial decrease in s4U content, ranging from 4.7-to 26-fold, was observed in tRNA samples from cells overexpressing RudS encoding genes (Figure 2A). Conversely, overexpression of stand-alone TudS or YbgA (stand-alone DUF1722 domain) encoding genes did not exhibit any statistically significant effect on the s4U content of total tRNA compared to control samples (Figure 2A).

**Figure 2.**
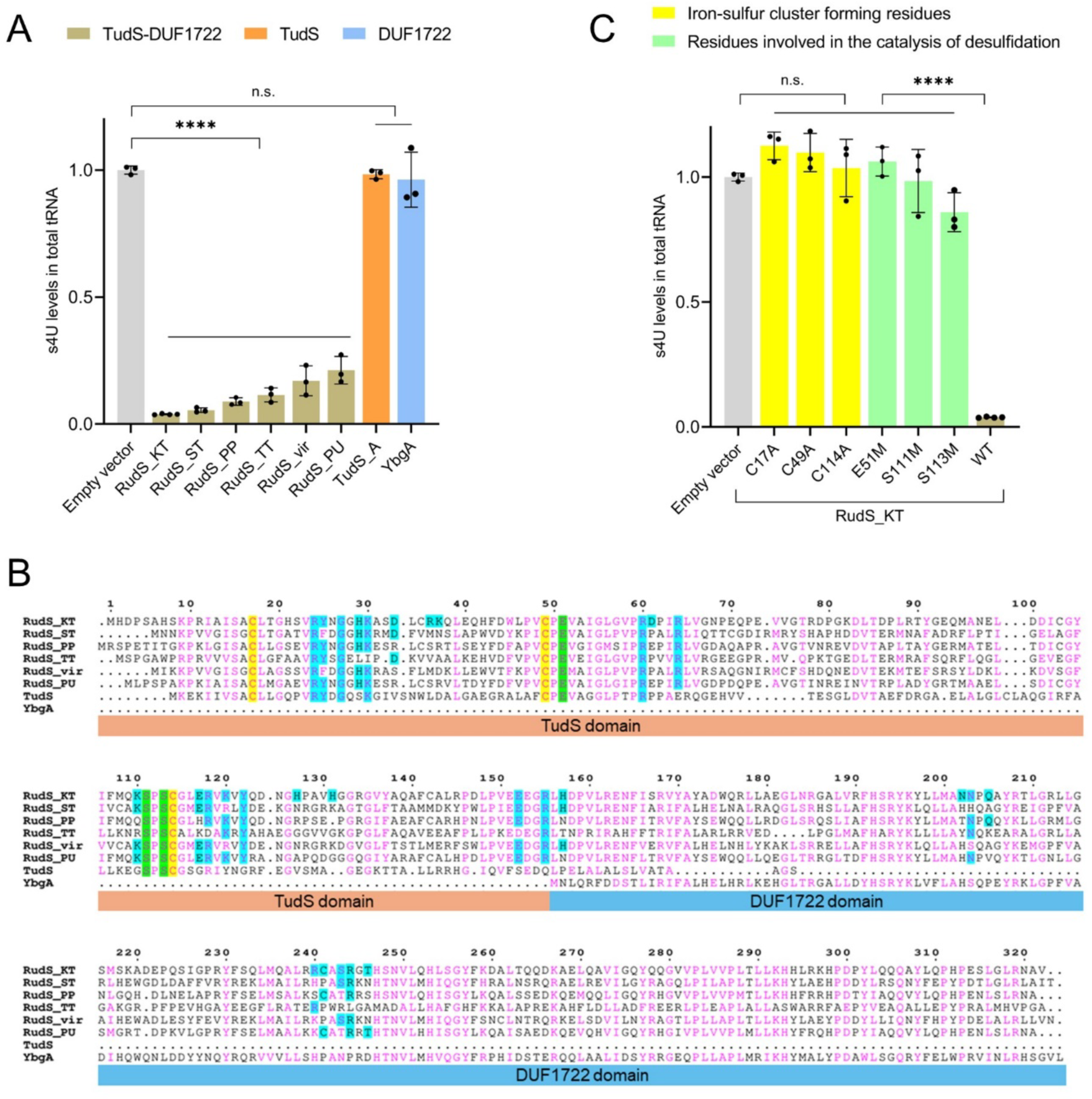
RudS reduces the s4U content of tRNA in vivo. A: s4U levels in *E. coli* total tRNA upon expression of RudS, TudS and DUF1722 encoding genes. ****: p<0.0001, n.s.: p≥0.05 compared to WT, one way ANOVA with Dunnet’s post hoc test. B: Sequence alignment of investigated RudS orthologs, TudS, and YbgA (stand-alone DUF1722 domain). Conserved cysteine residues involved in iron-sulfur cluster formation are highlighted in yellow. Conserved glutamic acid and serine residues catalyzing 4-thiouridine desulfidation reaction are highlighted in green. Amino acids predicted in tRNA binding and/or catalysis are highlighted in cyan. C: s4U levels in *E. coli* total tRNA upon expression of RudS_KT with single amino acid substitutions in conserved iron-sulfur cluster forming and catalytical residues. ****: p<0.0001, n.s.: p≥0.05 compared to WT, one way ANOVA with Dunnet’s post hoc test.

Among the six overexpressed RudS orthologs, the most significant decrease in s4U content (26-fold, compared to the negative control) was observed with RudS_KT from *Pseudomonas putida* KT2440, followed by RudS_ST from *Salmonella enterica* (18.3-fold), while RudS_PU from *Pseudomonas* sp. MIL9, overexpression resulted in the smallest decrease (4.7-fold). The variation in s4U content reduction among different RudS proteins raises the question of whether this was due to differences in heterologous expression levels or the specificity of RudS. To evaluate the levels of soluble proteins produced in *E. coli*, His-tags were introduced to the N-termini of RudS proteins for detection with anti-His-Tag antibodies.

Western blot analysis revealed that RudS_KT, while causing the most significant reduction in tRNA s4U content (Figure 2A), did not demonstrate the highest protein levels in the cells (Supplementary Figure S2A). In contrast, RudS_TT showed over 1.7-fold higher expression levels, but a lower s4U content reduction (8.7-fold, Figure 2A). The lower desulfidation activity of RudS_TT might be attributed to suboptimal temperature conditions for a protein originating from a thermophilic organism. It might also be due to inefficient substrate recognition, as tRNAs in thermophilic microorganisms often possess unique body modifications not found in their mesophilic counterparts (85). The detected soluble fractions of RudS_ST and RudS_PP were lower in comparison to RudS_KT (Supplementary Figure S2A), aligning with the observed reduction of s4U content (Figure 2A). Despite the undetectable level of soluble protein, a reduction in tRNA s4U content was observed with RudS_PU overexpression (Figure 2A). RudS_vir, on the other hand, stood out with relatively high protein levels (Supplementary Figure S2A), comparable to those of RudS_ST, yet its tRNA desulfidation activity appeared to be lower (Figure 2A). All tested RudS variants exhibited activity towards tRNA, indicating that RudS enzymes function as tRNA de-modifying erasers, targeting the 4-thiouridine modification in tRNA molecules.

### A functional 4Fe-4S cluster is needed for in vivo tRNA desulfidation by RudS

Previously we uncovered the mechanism of 4-thiouracil-containing compound desulfidation by TudS (41, 60). The key component in this process is a 4Fe-4S cluster harbored in the enzyme’s active center, bound by three cysteine residues, with the fourth iron atom engaging in substrate binding. Given that the overexpression of YbgA (the *E. coli* stand-alone DUF1722 domain) had no impact on tRNA s4U content (Figure 2A), we hypothesized that the TudS domain in the RudS fusion proteins should be responsible for the observed decrease of s4U in tRNA samples. We identified conserved cysteine residues (Figure 2B) which are likely to be involved in 4Fe-4S cluster formation (Table 1), created single amino acid substitutions of these cysteines in RudS homologues, and tested their tRNA desulfidation activity. Western blot analyses of His-tagged variants of mutant RudS proteins in the soluble fractions of the bacterial lysates indicated that the amounts of the mutant RudS protein variants are up to ∼4-fold lower than those of the wild types (Supplementary Figure S2B). The majority of mutant RudS protein variants showed a loss of activity (Figure 2C and Supplementary Figure S3), failing to significantly reduce s4U levels in *E. coli* tRNA, thus confirming our hypothesis. A single exception was the RudS_TT C47A protein variant (Supplementary Figure S3), which reduced the tRNA s4U content to levels comparable with those of the wild type RudS_TT. This suggests that a protein lacking one of the three cysteines might still form an active FeS cluster using only two cysteine ligands, a phenomenon previously reported in *Bacteroides thetaiotaomicron* fumarase. Despite lacking a typical third 4Fe-4S cluster-coordinating cysteine found in bacterial fumarases, this enzyme remains active but is more prone to oxidative stress (86).

### In vitro activity of RudS enzyme

Our previous studies showed that the iron-sulfur cluster in TudS is highly labile when exposed to oxygen, leading to an inactive enzyme when purified aerobically (40). Nevertheless, we produced N-terminally His-tagged RudS_KT in *E. coli* and purified it (Supplementary Figure S4A) under aerobic conditions with minimal oxygen exposure, as detailed in the methods section. On a molecular size exclusion chromatography column RudS_KT eluted as a 37.1 ± 0.7 kDa protein (theoretical molar mass of ≈39 kDa), indicating a monomeric architecture (Supplementary Figure S4B). The recombinant protein exhibited a brown color typical for [Fe-S] cluster containing proteins. Iron content determination using ferene assay indicated the presence of 1.01±0.13 Fe per protein monomer, suggesting the major population of protein having damaged Fe-S cluster assumably caused by the exposure to oxygen. Despite the poor iron content, UV spectrum of purified protein demonstrated spectrum characteristic for [4Fe-4S] clusters, which was partially bleached upon treatment with dithionite (Supplementary Figure S4C).

Despite indications of a damaged [4Fe-4S] cluster, purified RudS_KT remained soluble over extended periods of time. To test direct tRNA binding, as is typical for tRNA modifying enzymes, we conducted an electromobility shift assay (EMSA) with *E. coli* pseudouridine synthase (TruB) serving as a positive shift control (87). EMSA revealed a concentration-dependent protein band shift, confirming the RudS_KT-tRNA complex (Figure 3A).

**Figure 3.**
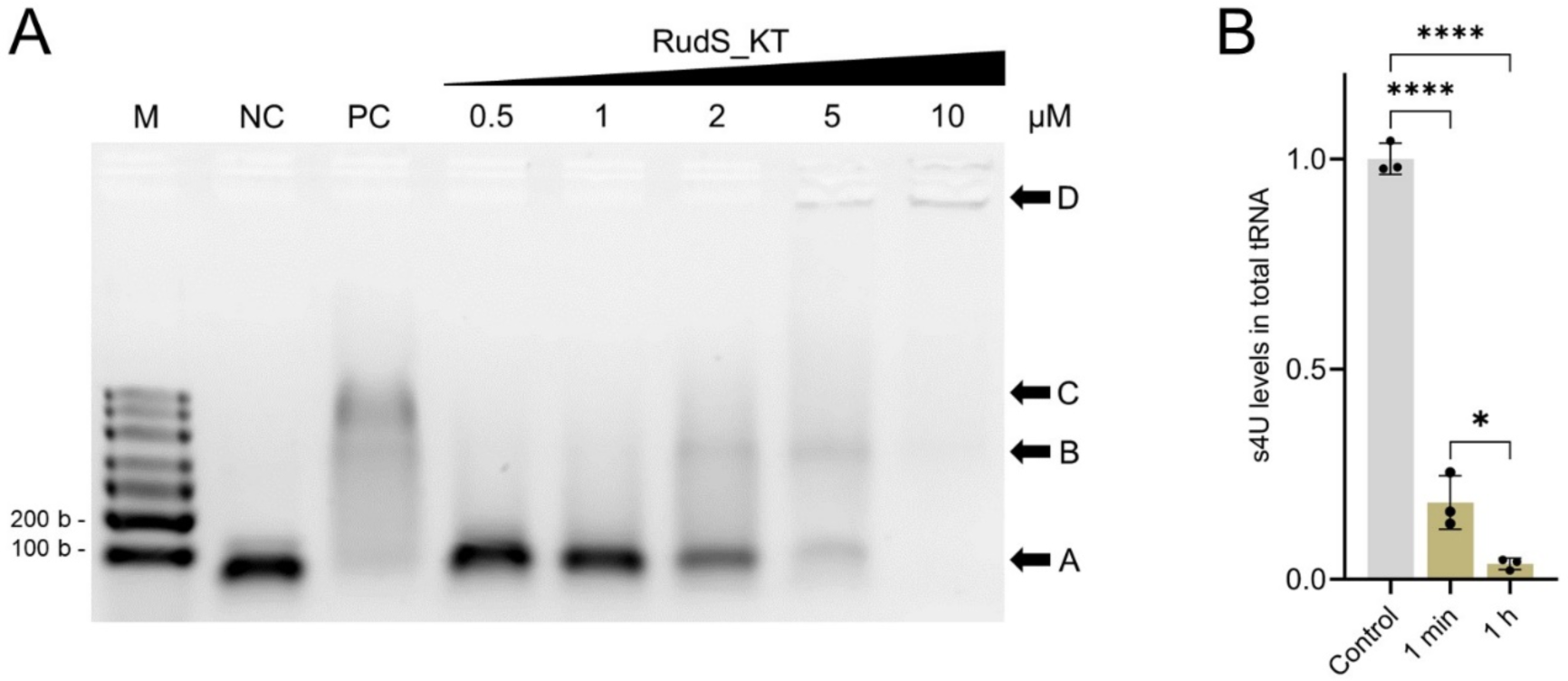
Purified recombinant RudS_KT interacts with tRNA in vitro. A: Electrophoretic mobility shift assay. M – molecular marker, NC – tRNA control, PC – positive shift control (1:5 molar ratio, using *E. coli* TruB pseudouridine synthase); 0.5–10 µM or RudS_KT was added to binding mixture resulting in 1:1–1:20 tRNA–RudS_KT molar ratios. Arrowhead A indicates unbound tRNA, B – tRNA-RudS_KT complex, C – tRNA-TruB complex, D – gel wells. B: Recombinant RudS_KT desulfidation activity in vitro using *E. coli* total tRNA. ****: p<0.0001, *: p<0.05, n.s.: p≥0.05, one way ANOVA with Bonferroni’s post hoc test.

To ensure that RudS_KT is directly responsible for the observed decrease in tRNA s4U content, we performed an in vitro tRNA desulfidation reaction using total *E. coli* tRNA. Contrary to previous observations with TudS, the aerobically purified and desalted RudS_KT protein retained activity. Approximately 13-fold molar excess of aerobically purified RudS_KT reduced the 4-thiouridine content in tRNA by ∼80% over 1 min, and by ∼95% over 1 h (Figure 3B). The desulfidation of the substrate was incomplete, likely due to the degradation of the remaining iron-sulfur clusters, although the inaccessibility of s4U due to non-physiological conditions or incorrect tRNA folding cannot be ruled out.

Additionally, we tested whether alarmones guanosine pentaphosphate and tetraphosphate influence RudS in vitro. We observed no statistically significant changes in the RudS enzymatic activity when using 1:10 concentration ratio of RudS to (p)ppGpp (Supplementary Figure S5).

### Docking studies reveal RudS amino acid residues involved in tRNA-binding

We were unable to successfully crystallize the RudS_KT protein and employed molecular docking and molecular dynamics (MD) simulations to explore its ability to desulfurize tRNAs. MD simulations of the holoenzyme were carried out as described in the Materials and Methods. The superposition of the top five models generated by both Alphafold2 and trRosetta is depicted in Figure 4A, indicating substantial similarity among the produced models. Figure 4B illustrates the modeling confidences reported by the respective methods, showing high confidence levels (plDDT>90). A structural comparison with the closest identified structural homologue (HHpred (88); PDBID:6z96) is presented in Figure 4C, revealing a resemblance in the organization of the entire TudS domain. The active center featuring the iron-sulfur cluster possesses identical amino acids, except for Lys98 in TudS being replaced with Met108 in RudS_KT. Hence, we propose that the catalytic mechanism of desulfuration could be analogous to the previously suggested mechanism (60). The model of the RudS_KT holoenzyme was constructed using the best model from Alphafold2 and transferring the iron-sulfur cluster from PDBID:6z96 (Fig. 4C) (60). The DUF1722 domain of RudS_KT comprises two compact groups of four alpha helices. No significant structural matches were identified using Dali (89) structural homology search for this segment of the model.

**Figure 4.**
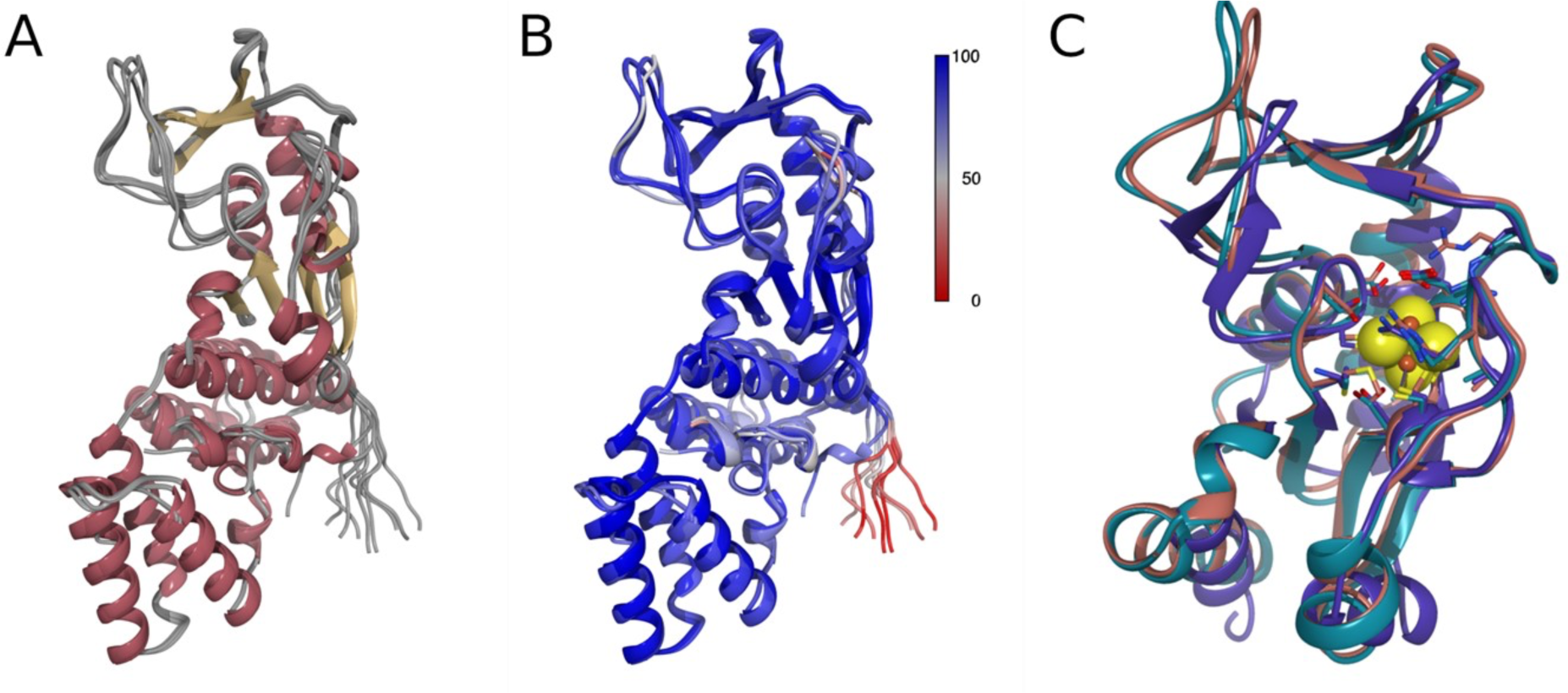
Structural comparison of RudS_KT models. A: a superposition of Alphafold2 and RosettaFold models colored by secondary structure. B: models colored by pLDDT. C: structural comparison of best Alphafold2 (teal) and trRosetta (salmon) models with closest homologue (PDBID:6z96) with structure identified with HHpred (purple). Iron-sulfur cluster represented with spheres).

Two independent 160 ns MD simulations of holoenzyme were carried out using randomly selected frames from restraint relaxation MD simulation’s trajectory. During simulations, the DUF1722 domain was mobile relative to the catalytic domain. Therefore, for RMSD and principal component analysis, we used only the compact part of Ruds_KT catalytic domain – amino acids from 11th to the 168th. Figure 5A illustrates RMSD changes during the simulation with the first structure as a reference. Essentially, the same ∼1.7 Å RMSD drift in both simulations from the reference indicated that the structure of the catalytic domain essentially converged. The PCA analysis in Figure 5B and C indicates the same conclusion, which is further supported by the visual inspection of the superposition in Figure 5D. In the latter one, the closest known structure of a homologous protein is added as well (TudS, PDBID:6z96). The structure of the catalytic domain stays essentially the same during all simulation time. The most important differences from TudS are an extended loop consisting of 116–135 amino acids (the same difference is evident in the Alphafold and Rosetta models superimposed with TudS in Figure 4) and a mobile loop consisting of 18–33 amino acids. We hypothesize that the mobility of these loops makes the active center of the enzyme more open and allows it to accommodate large substrates such as tRNAs at the cost of the ability to accommodate much smaller substrates such as thiouracil and similar ones.

**Figure 5.**
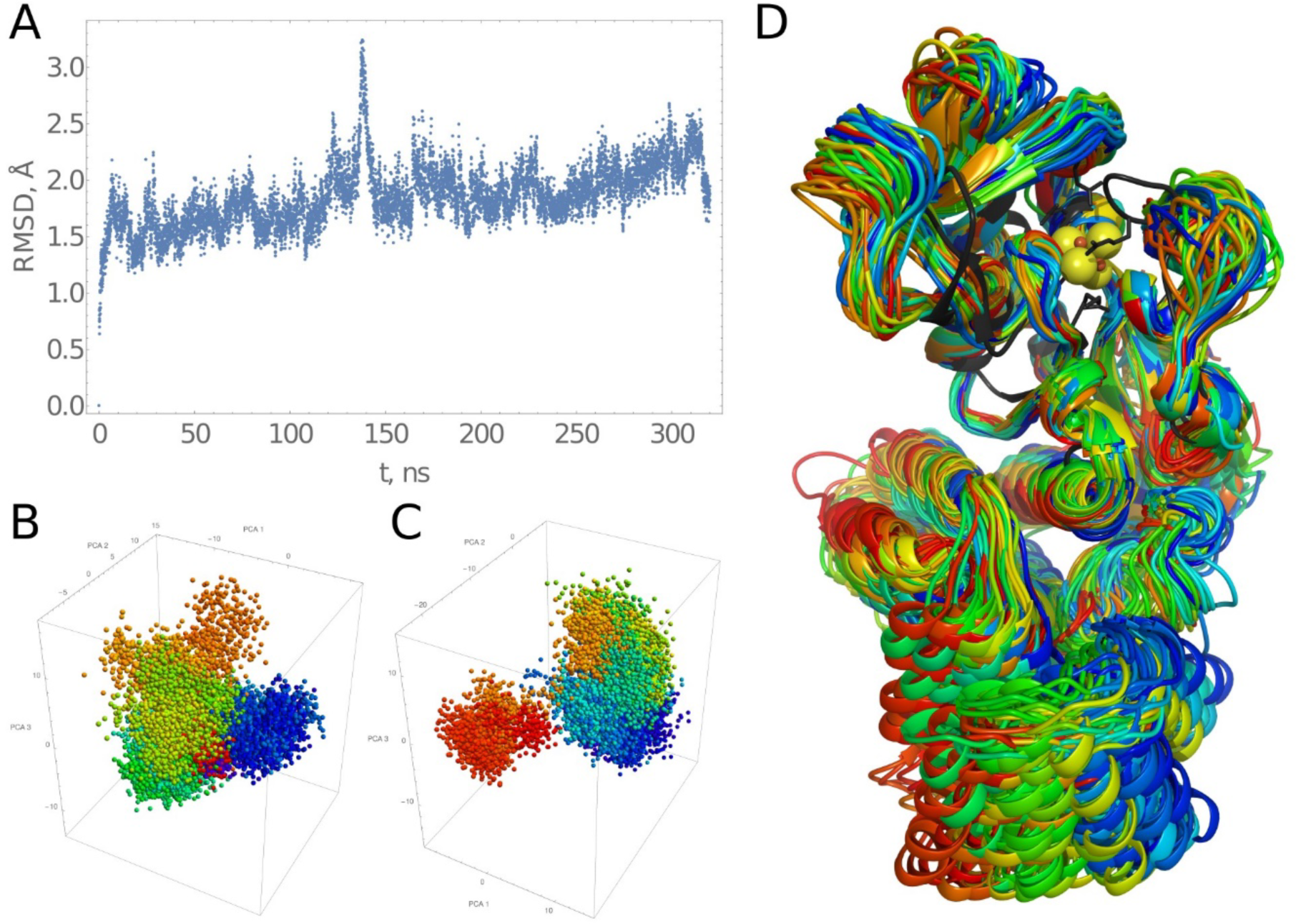
Molecular dynamics simulation of RudS_KT holoenzyme. A: cumulative RMSD plot of two 160 ns MD simulations with different starting conformations. B and C: PCA of both simulations. D: superposition of complexes colored according to the PCA, black – TudS protein (PDBID:6z96).

We carried out a series of molecular dynamics simulations for a selected tRNA structure as well. A phenylalanine tRNA from *E. coli* with a thiouridine modification at the 8th position was chosen as a model substrate for further docking studies (PDBID:6y3g). The buried thiouridine moiety in the structure was inaccessible for enzymatic attack. It is unlikely for the thiouridine to spontaneously flip out of the buried state during MD simulation; therefore, we manually flipped this residue out of the structure and carried out 500 ns MD simulations using the same methodology as employed for the holoenzyme, but without any restraints. The RMSD, PCA, and structural comparison reveal that the general ribose and phosphate backbone remained stable during the simulation (Supplementary Figure S6).

### Mutation analysis gives insight into tRNA-binding mechanism

The initial structures of the RudS holoenzyme and the tRNA with the thiouridine exposed were utilized for subsequent docking studies with HDOCK. Resultant complexes were ranked based on the distance between the relevant iron in the iron-sulfur cluster and the sulfur atom in the tRNA’s thiouridine. The top 15 complexes are depicted in Supplementary Figure S7. None of these complexes exhibited iron-sulfur distances relevant for actual catalysis. Nonetheless, these structures provided a preliminary hypothesis regarding how the tRNA might bind to the holoenzyme. Using these structures, we selected six groups of amino acids that are close to tRNA and might play a role in enzymatic catalysis and/or substrate binding (Supplementary Table 4).

We experimentally investigated these and hypothesized catalytic amino acids by mutating them into methionines (Table 2). Mutations of proposed catalytic amino acids essentially abolished desulfidase activities (Figure 2C, Supplementary Figure S3). Other mutations diminished or unaffected catalytic activities (Supplementary Figure S8). Each of the previously listed groups contains at least one amino acid affecting enzymatic activity. These experimental results were compared against molecular dynamics simulations of two holoenzyme-tRNA complex models (model 1 and model 2) obtained as described in the Supporting information.

**Table 2.** Effect of mutagenesis of predicted RudS_KT amino acids involved in enzymatic catalysis and/or substrate binding for s4U content in total *E. coli* tRNA. AVG – relative s4U content, stdev – standard deviation. Statistical significance was calculated by applying one way ANOVA with Dunnet’s post-hoc test. ****: p<0.0001, ***: p<0.001, **: p<0.01, n.s.: p≥0.05 compared to WT.

| Mutant | AVG | stdev | p-value | Group | Remarks |
| --- | --- | --- | --- | --- | --- |
| WT | 0.04 | 0 | - | - | Wild type RudS_KT |
| E51M | 1.06 | 0.06 | **** | CAT | Catalytic amino acid; conserved; removes proton from water molecule |
| S111M | 0.98 | 0.13 | **** | CAT | Catalytic amino acid; conserved; stabilizes Glu51 |
| S113M | 0.86 | 0.08 | **** | CAT | Catalytic amino acid; stabilizes Glu51 |
| R24M | 0.14 | 0.02 | ** | G1 | Interacts with Gly27 backbone carbonyl, Asn26 side chain carbonyl and nearby ribose ring oxygen. |
| Y25M | 0.07 | 0.03 | n.s. | G1 | Shields active center from the tRNAs phosphates. Opportunistically forms hydrogen bond with ribose OH group |
| Y25A | 0.17 | 0.01 | **** | G1 |  |
| Y25F | 0.04 | 0 | n.s. | G1 |  |
| N26M | 0.14 | 0.06 | ** | G1 | Interacts with Tyr25 forming hydrogen bond |
| G27M | 0.16 | 0.01 | *** | G1 | Part of flexible loop, forms hydrogen bond with Arg24 via backbone carbonyl |
| H29M | 0.07 | 0 | n.s. | G1 | No important interactions observed |
| K30M | 0.05 | 0.01 | n.s. | G1 | Occasionally interacts with tRNA phosphates and Asp33 |
| D33M | 0.04 | 0.01 | n.s. | G1 | No important interactions observed (salt bridge with Arg36) |
| R36M | 0.05 | 0.02 | n.s. | G1 | No important interactions observed (salt bridge with Asp33) |
| K37M | 0.06 | 0.03 | n.s. | G1 | No important interactions observed |
| R60M | 0.89 | 0.11 | **** | G2 | Catalytic amino acid; stabilizes Glu51 |
| D61M | 0.05 | 0.03 | n.s. | G2 | Occasionally interacts with bases in the tRNA |
| R64M | 0.15 | 0.05 | *** | G2 | Occasionally interacts with riboses in the tRNA |
| K110M | 0.05 | 0.02 | n.s. | G3 | Interacts with thiouridine base ribose and phosphate in tRNA |
| E117M | 0.06 | 0.02 | n.s. | G3 | Salt bridge with Arg155 |
| R118M | 0.09 | 0.01 | n.s. | G3 | Interaction with tRNAs riboses |
| K120M | 0.1 | 0.01 | n.s. | G3 | Interaction with tRNAs phosphates |
| Y122M | 0.1 | 0.01 | n.s. | G3 | Orientation of Lys120 and Arg64 |
| H127M | 0.04 | 0.01 | n.s. | G3 | No interaction |
| H131M | 0.06 | 0.02 | n.s. | G3 | No interaction |
| K110M,<br>K120M | 0.28 | 0.05 | **** | G3 | Synergistic effect, Lys110 partial compensation by Lys120 |
| E152M | 0.15 | 0.01 | *** | G4 | Salt bridge with Arg155 |
| R155M | 0.24 | 0.02 | **** | G4 | Salt bridge with Glu152, Van der Waals interaction with tRNA ribose |
| H157M | 0.09 | 0 | n.s. | G4 | Hydrogen bond with ribose ring oxygen |
| N202M | 0.21 | 0 | **** | G5 | Backbone stabilization via side chain in turn |
| N203M | 0.36 | 0.06 | **** | G5 | Backbone stabilization via side chain in turn |
| Q205M | 0.04 | 0.01 | n.s. | G5 | No interaction |
| R240M | 0.06 | 0.02 | n.s. | G6 | No interaction |
| C241M | 0.16 | 0.04 | **** | G6 | No interaction |
| S243M | 0.47 | 0.06 | **** | G6 | Interaction with anticodon base |
| R244M | 0.53 | 0.07 | **** | G6 | Interaction with phosphate in anticodon region |
| T246M | 0.18 | 0.02 | **** | G6 | Backbone interaction via side chain |

The superposition of catalytic domains (from 11th to 168th amino acids) for these models and the holoenzyme is illustrated in Figure 6A. These models are very similar except previously discussed mobile loops, which in the case of the enzyme-substrate complex are somewhat less mobile. Both models allow us to propose that the DUF1722 domain interacts with the anticodon arm and anticodon of the tRNA substrate, whereas the catalytic domain interacts with the variable loop, acceptor stem, and D arm of tRNA. In both models of the enzyme-substrate complex, mobility of the DUF1722 domain relative to the catalytic one is evident. This mobility, probably, is needed to accommodate a range of tRNA substrates with thiouridine modification. The RMSD of the catalytic domain is stable during the last 100 ns for model 2 (only thiouridine S – iron-sulfur cluster Fe restraint present). The structure revisits catalytic relevant conformations where Ser113 hydroxy hydrogen – Glu51 sidechain carboxy oxygen distances reach ∼2 angstroms over entire 260 ns simulation length.

**Figure 6.**
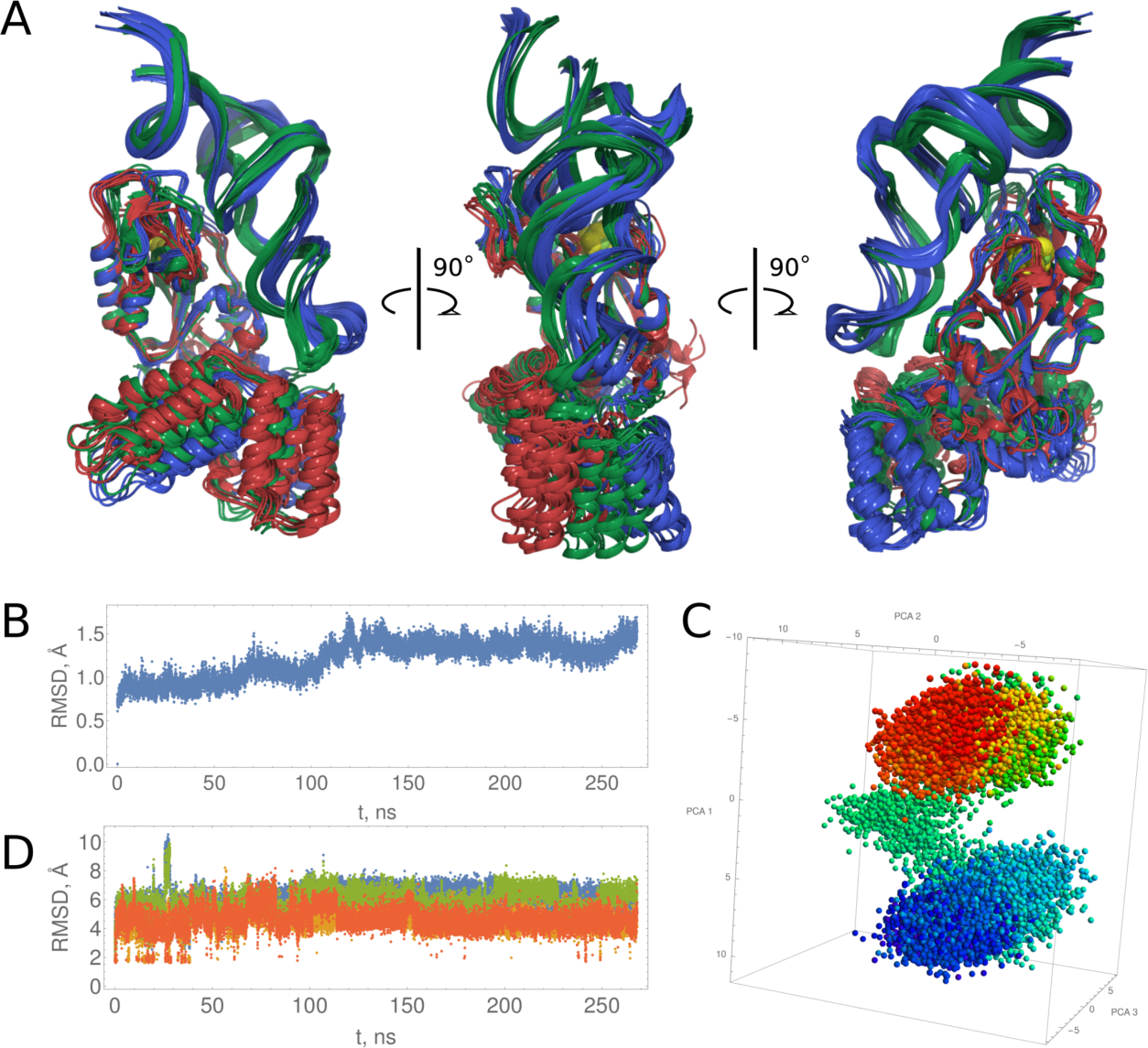
A: Superposition of model 1 (blue), model 2 (green), and homoenzyme (red) final models. B: Model 2 RMSD of catalytic domain. C: model 2 PCA of catalytic domain. D: Red and orange Ser113 hydroxy hydrogen – Glu51 sidechain carboxy oxygen distances, green and blue Ser111 hydroxy hydrogen – Glu51 sidechain carboxy oxygen distances.

Both models feature an additional base participating in tRNA-enzyme active site binding (Figure 7A). The base U45 (PDBID:6y3g) is not paired in the experimental tRNA structure and was dangling outside during initial tRNA simulations both with thiouridine side chain paired inside and manually flipped outside. To our best knowledge, homologous flipped uridine is present in at least three more known *E. coli* tRNA structures – valine tRNA (pdbid:7eqj), initiator formylmethionine tRNA (pdbid:5l4o), and *E. coli* aspartate tRNA (pdbib:6ugg). The uridine participates in an extensive network and forms a double hydrogen bond with Asn26. Nearby residing Arg24 suggests the possibility of a cation-π stabilizing interaction. Arg24 is oriented by a hydrogen bond with Gly27 backbone carbonyl oxygen atom. These structural observations apply for both models. Mutation of these amino acids to methionine results in diminished RudS_KT enzymatic desulfidase activity. A somewhat smaller change than anticipated in enzymatic activity is explainable by the methionine’s ability to form hydrogen bonds. Tyr25 mutation to methionine resulted in mutant protein functionally identical to the WT. We constructed two additional mutants: Tyr25 to alanine, which had decreased activity, and Tyr25 to phenylalanine, which had the same activity as WT.

**Figure 7.**
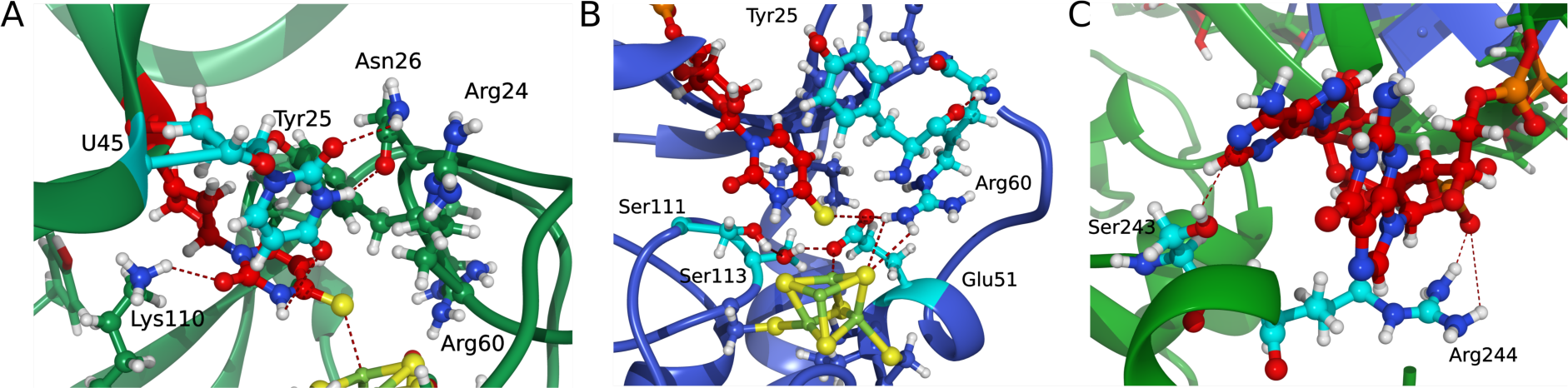
A: tRNA-RudS_KT active site binding. B: The active center of model 1 with four restrained catalytic distances. C: Ser243 and Arg244 participate in the interaction with anticodon bases and anticodon phosphate backbone.

The amino acids previously deduced from RudS homology with TudS as catalytic ones (Glu51, Ser111, and Ser113) produced inactive enzymes when mutated to methionines (Figure 2C). We identified an additional catalytic amino acid, Arg60, which function is both to orient Glu51 and provide a positive charge, which could additionally stabilize transitional states of proton removal from water and nucleophilic attack of thiouridine with produced hydroxide ion. This arginine is fully conserved in the entire family of homologous proteins, and retrospectively, we identified a homologous arginine serving the same purpose in the experimentally determined structure of TudS (60). The comparison of randomly selected frames from model 2 and model 1 molecular dynamics shows that orientations of catalytic amino acids relative to the thiouridine moiety are very similar (Supplementary figure S9) in both models. Therefore, for clearer representation of the active center, we chose to illustrate the active center of model 1 (model with four restrained catalytic distances) (Figure 7B).

The DUF1722 domain in RudS_KT interacts with both the anticodon stem and the anticodon of tRNAs. While the molecular dynamics simulations may not depict the final binding state accurately, specific frames identified shed light on the significance of Ser243 and Arg244. Our hypothesis suggests that these amino acids, potentially in conjunction with other unidentified amino acids, play a role in interacting with the anticodon bases and the anticodon phosphate backbone (Figure 7C). The summary of the rationale behind the enzymatic activities of other mutants is briefly outlined in Table 2. These models serve as explanatory tools for most of the experimentally observed desulfidase activities of the mutants, although they should be regarded as hypotheses rather than definitive structural representations.

## DISCUSSION

While the understanding of TudS desulfidases is evolving, the function and structure of the Domain of Unknown Function 1722 had remained elusive until now. Previous studies from our group had suggested that additional domains fused to TudS might influence its substrate preferences, potentially even accommodating intact tRNA molecules (40, 41, 60). This study successfully demonstrates that the DUF1722 domain facilitates the targeted removal of s4U modification from tRNA molecules by the RudS (TudS-DUF1722 fusion) proteins both in vivo and in vitro.

### Structural Insights and Catalytic Mechanism

Structural examination of RudS_KT from *Pseudomonas putida* KT2440 was conducted using molecular docking and molecular dynamics simulations, with subsequent experimental investigation connecting the proposed models and experimentally observed desulfidase activities of mutants. The investigation of the hypothesized catalytic amino acids (Glu51, Ser111, Ser113, and Arg60) and cysteines holding iron-sulfur cluster (Cys17, Cys49 and Cys114) by mutational analysis essentially abolished the RudS desulfidase activity, confirming their critical role in catalysis. Additionally, the models identified other important residues (Asn26, Arg24, Tyr25, Ser243, Arg244) for tRNA binding and catalysis, and their mutations resulted in diminished RudS_KT enzymatic activities. The effects of single amino acid substitutions to methionine (63) were all in line with the models we propose, except in the case of Tyr25, where two additional mutants were employed. The Tyr25 mutation to methionine was functionally identical to WT enzyme, which is at odds with its position in the modeled complex – Tyr25 provides shielding of the active center from tRNA’s phosphates and their negative charge, which would destabilize the transitional state and disrupt catalysis. The RudS_KT Tyr25 to alanine mutant displayed decreased enzymatic activity, while the Tyr25 to phenylalanine mutant retained the same activity as WT. We propose that methionine is large enough to provide substantial electrostatic shielding; therefore, only the Tyr25 to alanine variant function was significantly affected by the substitution.

The proposed models of the RudS enzyme-tRNA substrate complex suggest that the DUF1722 domain primarily interacts with the anticodon arm and anticodon of the tRNA substrate, while the catalytic TudS domain interacts with the variable loop, acceptor stem, and D arm of the tRNA. Moreover, the elongated loops encompassing amino acids 116-135 and 18-33, along with the mobility of the DUF1722 domain, likely aid in the recognition and adaptation of a diverse array of tRNAs containing the 4-thiouridine modification at position 8. This functional specialization between the two domains likely enables the precise elimination of the s4U modification from the tRNA molecule by RudS.

### Evolutionary Insights and Functional Implications

The DUF1722 genes are predominantly reported in bacteria (94.2% of all sequences) with archaeal DUF1722 making up only 5.1%. 74.4% of DUF1722s are fused with the TudS gene and only 21.4% of known DUF1722 genes encode a stand-alone protein (81). Genes encoding RudS are often found in light-responsive operons (Supplementary Figure S10), which are responsible for UV-induced DNA damage repair (i.e., DNA-photolyases) or light-induced oxidative stress response (i.e., carotenoid biosynthesis gene clusters) (90). Recent studies in the genus *Pseudomonas* have revealed the role of light-inducible transcriptional regulators in controlling the expression of light-responsive genes. These regulators belong to the MerR family and are adenosyl B12-dependent, making them sensitive to light (class II LitR regulators). In the dark, LitR regulators function as negative regulators, suppressing the transcription of light-inducible genes. However, when exposed to 450 nm blue light, LitRs become deactivated. One of the operons controlled by this mechanism encodes PhrB DNA photolyase, LitR, and RudS (91, 92).

At the same time, the tRNA s4U modification is known to act as a photosensitive residue in tRNA, crosslinking with neighboring cytidine in 13th position upon exposure to near-UV radiation. Near-UV light triggers a bacterial growth delay effect causing some tRNA species to become poor substrates for aminoacylation resulting in accumulation of uncharged tRNA and transient cessation of protein synthesis (30, 93). The accumulation of ribosomes stalled with non-aminoacylated tRNA are known to initiate a stringent response (94), which is the most likely mechanism causing alarmone guanosine tetraphosphate (ppGpp) accumulation after cell irradiation by UV-A (95, 96). Paradoxically, *E. coli* Δ*thiI* mutants, incapable of s4U tRNA modification biosynthesis, are less susceptible to UVA, as s4U is the primary target of this irradiation. Subsequently, this eliminates a damaging synergistic effect of UVA and UVB irradiation observed in *E. coli* (97). On the other hand, studies with *Salmonella typhimurium* revealed that 4-thiouridine plays a crucial role in resistance to near-UV irradiation, with mutants lacking 4-thiouridine and those deficient in ppGpp synthesis being sensitive to near-UV-induced killing; it further suggested a model wherein ppGpp and ApppGpp induce the synthesis of a set of then unidentified proteins essential for resistance to near-UV irradiation in response to the cross-linking of 4-thiouridine in tRNA (96). It is worth noting that *Salmonella typhimurium* possesses a RudS encoding gene (RudS_ST in this study, ORF319 previously) under control of transcription regulator of MerR family (83), while *E. coli* does not.

The prevalence of DUF1722 genes in bacteria, often fused with TudS, and their association with light-responsive genes, suggest a light-induced mechanism for prokaryotic tRNA s4U de-modification. This mechanism may play a role in regulating alarmone levels during UV-induced tRNA cross-linking and the stringent response, serving as a bacterial survival strategy under UV stress. Although in vitro experiments showed no changes in the enzymatic activity of RudS_KT in the presence of the (p)ppGpp molecules, this does not rule out the possibility that the alarmones and RudS are part of the same regulatory mechanism within the cell, interacting through currently unknown intermediary factors or pathways.

This hypothesis is further supported by the case of *Enterobacter cloacae*, which demonstrated that pre-treating bacteria with sub-lethal doses of UVA radiation activates an unidentified mechanism, leading to a reduction in the 4-thiouridine content of tRNA that helps to evade the growth delay caused by crosslinked tRNA during subsequent UVA irradiation (98). In the genome sequence of *E. cloacae* (GenBank: CP135498.1), the RudS encoding gene (locus tag: RRL13_00330) is in the immediate vicinity to MerR family transcriptional regulator (locus tag: RRL13_00340), suggesting a light inducible RudS gene expression, which might explain the observed effect. Furthermore, a previous study by the same authors revealed that after the exposure to UVA, not only a growth delay was induced in both *E. cloacae* and *E. coli*, but also a burst of ppGpp was observed (95). However, the ppGpp amounts accumulated in *E. coli* were twice those reached in *E. cloacae*. Moreover, the time needed to restore ppGpp content to basal levels in *E. cloacae* was shorter than that required in *E. coli*. This period aligned with the time when growth resumed at its normal rate in both species. It is worth noting that *E. coli* genome encodes a stand-alone DUF1722 domain YbgA, while *E. cloacae* encodes a RudS fusion protein. Both these species lack a stand-alone TudS. We speculate that *E. cloacae*, which possesses a RudS fusion protein, was able to reduce the 4-thiouridine content of tRNA and mitigate the growth delay caused by cross-linked tRNA during UVA irradiation. In contrast, the *E. coli*, which lacks a stand-alone RudS and instead encodes a stand-alone DUF1722 domain (YbgA), exhibited a more pronounced stringent response and delayed recovery after UV exposure. These observations confirm that the function of RudS is different from that of a stand-alone DUF1722 and that of a stand-alone TudS. There is a possibility that during evolution, in certain species, the gene encoding RudS may have undergone a split, resulting in separate stand-alone TudS and stand-alone DUF1722 encoding genes.

### Conclusions

This study provides compelling evidence that RudS enzymes, consisting of a DUF1722 domain fused with the TudS desulfidase, facilitate the targeted removal of the s4U modification from tRNA molecules. We may speculate that certain bacterial species have evolved a light-inducible mechanism for tRNA s4U de-modification, which likely plays a role in regulating alarmone levels during a tRNA UV-crosslinking-induced stringent response-like state. By diminishing the available pool of tRNA substrate, such adaptive process might mitigate continuous cross-linking, thereby preventing translation derangement. Such strategic adjustment may serve as a survival mechanism for bacteria, particularly in the face of intense UV radiation exposure, ensuring the seamless operation of stress-response machinery.

## DATA AVAILABILITY

The models are available in ModelArchive at https://www.modelarchive.org/doi/10.5452/ma-6xegk.

## SUPPLEMENTARY DATA

Supplementary Data are available at NAR online.

## AUTHOR CONTRIBUTIONS

R. J.: Formal analysis, Investigation, Methodology, Validation, Visualization, Writing – original draft, Writing – review & editing. A. L.: Formal analysis, Methodology, Validation, Visualization, Writing – original draft, Writing – review & editing. D. P.: Investigation, Validation, Writing – review & editing. R. M.: Conceptualization, Funding acquisition, Resources, Writing – review & editing. A. A.: Conceptualization, Formal analysis, Investigation, Validation, Supervision, Visualization, Writing – original draft, Writing – review & editing.

## ACKNOWLEDGEMENTS

We thank Justas Vaitekūnas for technical assistance.

## FUNDING

This work was supported by the European Regional Development Fund under grant agreement with the Research Council of Lithuania (LMTLT) [01.2.2-LMT-K-718-03-0082 to RM].

## CONFLICT OF INTEREST

The authors declare no conflicts of interests.

## Notes

https://www.modelarchive.org/doi/10.5452/ma-6xegk

### Competing Interest Statement

The authors have declared no competing interest.

