## Supplementary Data for "RudS: Bacterial Desulfidase Responsible for tRNA 4-Thiouridine De-modification"

### SUPPORTING INFORMATION

### MATERIAL AND METHODS

**Supplementary Table 1.** Bacterial strains used in this study.

| Bacterial strain | Comments | Source |
|---|---|---|
| *Escherichia coli* DH5α | Used for routine DNA manipulations | Pharmacia, USA |
| *Escherichia coli* BL21(DE3) | Used for recombinant protein overexpression | Avidis, France |
| *Pseudomonas sp.* MIL9 | NCBI accession ID: PRJNA698458 | Isolated from soil; Vilnius university Life Sciences Center (1) |
| *Pseudomonas* sp. MIL19 | NCBI accession ID: PRJNA877084 | Isolated from soil; Vilnius university Life Sciences Center |
| *Pseudomonas putida* KT2440 | DSM No.: 6125 | Leibniz Institute DSMZ-German Collection of Microorganisms and Cell Cultures GmbH |
| *Salmonella enterica* subsp. *enterica* serovar Typhimurium LT2 Δ*pyrF* | Uracil auxotroph | Kind gift from Gunilla Jäger, Umeå University, Sweden |
| *Thermus thermophilus* HB8 | DSM No.: 579 | Leibniz Institute DSMZ-German Collection of Microorganisms and Cell Cultures GmbH |

**Supplementary Table 2.** Plasmid vectors used in this study.

| Plasmid vector | Comments | Source |
|---|---|---|
| pLATE11 | Empty vector constructed by amplification of backbone and blunt-end ligation. | Thermo Fisher Scientific, USA and this study |
| pLATE11-TudS | TudS encoding expression vector | (2) |
| pLATE11-YbgA | DUF1722 encoding expression vector | This study |
| pBR322-RudS_vir | Synthetic vector encoding viral RudS | Synthesized by Thermo Fisher Scientific, USA |
| pLATE11-RudS_KT | RudS encoding expression vectors | This study |
| pLATE11-RudS_ST | | |
| pLATE11-RudS_PP | | |
| pLATE11-RudS_TT | | |
| pLATE11-RudS_vir | | |
| pLATE11-RudS_PU | | |
| pLATE52-RudS_KT | RudS with N-terminal 6xHis-tag encoding expression vectors | This study |
| pLATE52-RudS_ST | | |
| pLATE52-RudS_PP | | |
| pLATE52-RudS_TT | | |
| pLATE52-RudS_vir | | |
| pLATE52-RudS_PU | | |
| pLATE52-YbgA | DUF1722 with N-terminal 6xHis-tag encoding expression vectors | This study |
| pLATE11-RudS_KT_C114A | RudS with conservative cysteine substitution encoding expression vectors | This study |
| pLATE11-RudS_KT_C49A | | |
| pLATE11-RudS_KT_C17A | | |
| pLATE11-RudS_ST_C12A | | |
| pLATE11-RudS_ST_C44A | | |
| pLATE11-RudS_ST_C110A | | |
| pLATE11-RudS_PP_C18A | | |
| pLATE11-RudS_PP_C50A | | |
| pLATE11-RudS_PP_C115A | | |
| pLATE11-RudS_TT_C16A | | |
| pLATE11-RudS_TT_C47A | | |
| pLATE11-RudS_TT_C111A | | |
| pLATE11-RudS_vir_C12A | | |
| pLATE11-RudS_vir_C44A | | |
| pLATE11-RudS_vir_C110A | | |
| pLATE11-RudS_PU_C15A | | |
| pLATE11-RudS_PU_C47A | | |
| pLATE11-RudS_PU_C112A | | |
| pLATE52-RudS_KT_C114A | RudS_KT with conservative cysteine substitution and N-terminal 6xHis-tag encoding expression vectors | This study |
| pLATE52-RudS_KT_C49A | | |
| pLATE52-RudS_KT_C17A | | |
| pLATE11-RudS_KT_R24M | RudS_KT with amino acid predicted to participate in substrate binding and/or catalysis substitution and encoding expression vectors | This study |
| pLATE11-RudS_KT_Y25M | | |
| pLATE11-RudS_KT_Y25A | | |
| pLATE11-RudS_KT_Y25F | | |
| pLATE11-RudS_KT_N26M | | |

| | | |
|---|---|---|
| pLATE11-RudS_KT_G27M | | |
| pLATE11-RudS_KT_H29M | | |
| pLATE11-RudS_KT_K30M | | |
| pLATE11-RudS_KT_D33M | | |
| pLATE11-RudS_KT_R36M | | |
| pLATE11-RudS_KT_K37M | | |
| pLATE11-RudS_KT_E51M | | |
| pLATE11-RudS_KT_R60M | | |
| pLATE11-RudS_KT_D61M | | |
| pLATE11-RudS_KT_R64M | | |
| pLATE11-RudS_KT_K110M | | |
| pLATE11-RudS_KT_S111M | | |
| pLATE11-RudS_KT_S113M | | |
| pLATE11-RudS_KT_E117M | | |
| pLATE11-RudS_KT_R118M | | |
| pLATE11-RudS_KT_K120M | | |
| pLATE11-RudS_KT_Y122M | | |
| pLATE11-RudS_KT_H127M | | |
| pLATE11-RudS_KT_H131M | | |
| pLATE11-RudS_KT_E152M | | |
| pLATE11-RudS_KT_R155M | | |
| pLATE11-RudS_KT_H157M | | |
| pLATE11-RudS_KT_N202M | | |
| pLATE11-RudS_KT_N203M | | |
| pLATE11-RudS_KT_Q205M | | |
| pLATE11-RudS_KT_R240M | | |
| pLATE11-RudS_KT_C241M | | |
| pLATE11-RudS_KT_S243M | | |
| pLATE11-RudS_KT_R244M | | |
| pLATE11-RudS_KT_T246M | | |
| pLATE52-RudS_KT_R24M<br>pLATE52-RudS_KT_N26M<br>pLATE52-RudS_KT_G27M<br>pLATE52-RudS_KT_H29M<br>pLATE52-RudS_KT_K30M<br>pLATE52-RudS_KT_E51M<br>pLATE52-RudS_KT_R60M<br>pLATE52-RudS_KT_R64M<br>pLATE52-RudS_KT_K110M<br>pLATE52-RudS_KT_S111M<br>pLATE52-RudS_KT_S113M<br>pLATE52-RudS_KT_R118M<br>pLATE52-RudS_KT_K120M<br>pLATE52-RudS_KT_Y122M<br>pLATE52-RudS_KT_E152M<br>pLATE52-RudS_KT_R155M<br>pLATE52-RudS_KT_H157M<br>pLATE52-RudS_KT_N202M<br>pLATE52-RudS_KT_N203M<br>pLATE52-RudS_KT_R240M | RudS with amino acid predicted to participate in substrate binding and/or catalysis substitution and N-terminal 6xHis-tag encoding expression vectors | This study |

| |
|---|
| pLATE52-RudS_KT_C241M |
| pLATE52-RudS_KT_S243M |
| pLATE52-RudS_KT_R244M |
| pLATE52-RudS_KT_T246M |

**Supplementary Table 3.** Oligonucleotide primers used in this study.

| Primer name | Sequence (>3') | Purpose |
|---|---|---|
| pLATE11_empty_FW | TGACTTCCCATCTCCGGTTT | Amplification of vector backbone for empty vector creation. |
| pLATE11_empty_RV | AGTTATATCTCCTTCTGGATTTAAATGTTA | |
| RudS_KT_52_FW | GGTTGGGAATTGCAACACGACCCTTCCGCCCAC | Amplification of *rudS_KT* gene for cloning into pLATE11 and pLATE52 vectors |
| RudS_KT _11_FW | AGAAGGAGATATAACTATGCACGACCCTTCCGCC | |
| RudS_KT _11_52_RV | GGAGATGGGAAGTCATCAGACCGCATTGCGCAAG | |
| RudS_ST_11_31_FW | AGAAGGAGATATAACTATGAATAACAAACCCGTAGTCGGC | Amplification of *rudS_ST* gene for cloning into pLATE11 and pLATE52 vectors |
| RudS_ST_11_52_RV | GGAGATGGGAAGTCATTAGGTGATGGCTAAGCGCAG | |
| RudS_ST_52_FW | GGTTGGGAATTGCAAAATAACAAACCCGTAGTCGGC | |
| RudS_PP_11_31_FW | AGAAGGAGATATAACTATGCGCTCACCAGAAACCATCACC | Amplification of *rudS_PP* gene for cloning into pLATE11 and pLATE52 vectors |
| RudS_PP _11_52_RV | GGAGATGGGAAGTCATTATAGGGCGTTACGCAAGCTG | |
| RudS_PP _52_FW | GGTTGGGAATTGCAACGCTCACCAGAAACCATCACC | |
| RudS_TT_11_31_FW | AGAAGGAGATATAACTATGAGCCCAGGGGCGTGG | Amplification of *rudS_TT* gene for cloning into pLATE11 and pLATE52 vectors |
| RudS_TT_52_FW | GGTTGGGAATTGCAAAGCCCAGGGGCGTGG | |
| RudS_TT_11_52_RV | GGAGATGGGAAGTCATTAGGCCCCGGGGACGTG | |
| RudS_PU_11_31_FW | AGAAGGAGATATAACTATGCTCCCGTCCCCTGCCAAAC | Amplification of *rudS_PU* gene for cloning into pLATE11 and pLATE52 vectors |
| RudS_PU_11_52_RV | GGAGATGGGAAGTCATTAGATGGCGTTTCGCAGGCTG | |
| RudS_PU_52_FW | GGTTGGGAATTGCAACTCCCGTCCCCTGCCAAAC | |
| RudS_vir_11_52_RV | GGAGATGGGAAGTCATTAGTTCACCAAGAGTCTTAGAGCCAGTT CATC | Amplification of *rudS_vir* gene for cloning into pLATE11 and pLATE52 vectors |
| RudS_vir_11_31_FW | AGAAGGAGATATAACTATGATAAAAAAACCTGTCGTTGGGATCA GTGG | |
| RudS_vir_52_FW | GGTTGGGAATTGCAAATAAAAAAACCTGTCGTTGGGATCAGTGG | |
| YbgA_11_31_FW | AGAAGGAGATATAACTATGAATCTACAACGATTTGATGACAG | Amplification of *ybgA* gene for cloning into pLATE11 and pLATE52 vectors. |
| YbgA_11_52_RV | GGAGATGGGAAGTCATTATAAAACTCCTGAATGGCGC | |
| YbgA_52_FW | GGTTGGGAATTGCAAAATCTACAACGATTTGATGACAG | |

| | | |
|---|---|---|
| RudS_ST_C12A | AGTCGGCATTAGCGGAGCTCTGACTGGCGCC | Site directed mutagenesis of conservative cysteines using pLATE11 or pLATE52 vectors with insert as a template. |
| RudS_ST_C44A | CCTACAAACCGATCGCCCCGGAGGTGGCGA | |
| RudS_ST_C110A | TGTGTGTGCAAAATCGCCCAGTGCCGGAATGGAGCG | |
| RudS_TT_C16A | TGTGGTGAGCGCCGCCCTGGGGTTCGCC | |
| RudS_TT_C47A | CTTCGTCCCCGTCGCCCCGGAGGTGGAG | |
| RudS_TT_C111A | CCGCTCCCCCTCCGCCGCCCTGAAGGAC | |
| RudS_KT_C17A | CGCCATCAGCGCCGCCCTGACCGGGCAC | |
| RudS_KT_C49A | GACTGGCTACCCGTGGCTCCGGAAGTGGCAAT | |
| RudS_KT_C114A | TCTTCATGCAGAAGTCACCTTCAGCCGGCCTGGAACG | |
| RudS_PP_C18A | TGGGCATCAGTGCCGCCTTACTGGGCTCCG | |
| RudS_PP_C50A | TTTCGCCCCCGTGGCCCCCGAAGTAGGG | |
| RudS_PP_C115A | CAGTCCCCGTCCGCCGGCCTGCACCG | |
| RudS_PU_C15A | CGCCATCAGCGCCGCCCTGATGGGCGCA | |
| RudS_PU_C47A | TTTCGTGCCGGTCGCCCCGGAAGTCGCG | |
| RudS_PU_C112A | GAAATCGCCGTCCGCCGGACTGGAGCGG | |
| RudS_vir_C12A | TCGTTGGGATCAGTGGTGCTTTGGCCGGTTCTTCTG | |
| RudS_vir_C44A | TGGAATGGGTAACATTCAAACCAGTAGCTCCGGAAATGGCTA | |
| RudS_vir_C110A | GTGCAAAATCTCCTAGCGCTGGTATGGAACGCGTGC | |
| RudS_KT_R24M | ACCGGGCACAGCGTGATGTACAACGGCGGCCAC | Site directed mutagenesis of residues predicted to participate in catalysis or/and substrate binding using pLATE11 or pLATE52 vectors with insert as a template. |
| RudS_KT_Y25A | GTGGCCGCCGTTGGCGCGCACGCTGTGC | |
| RudS_KT_Y25F | GGCCGCCGTTGAAGCGCACGCTG | |
| RudS_KT_Y25M | CTTGTGGCCGCCGTTCATGCGCACGCTGTGCCC | |
| RudS_KT_N26M | GTGGCCGCCCATGTAGCGCACGCTGTGC | |
| RudS_KT_G27M | GGAGGCCTTGTGGCCCATGTTGTAGCGCACGCT | |
| RudS_KT_H29M | CAGGTCGGAGGCCTTCATGCCGCCGTTGTAGCG | |
| RudS_KT_K30M | CGGAGGCCATGTGGCCGCCGTTGT | |
| RudS_KT_D33M | GCTGTTTACGGCACAGCATGGAGGCCTTGTGGCCG | |
| RudS_KT_R36M | AAGGCCTCCGACCTGTGCATGAAACAGCTGGAACAGCAC | |
| RudS_KT_K37M | GTGCTGTTCCAGCTGCATACGGCACAGGTCGGAG | |
| RudS_KT_E51M | CCAAGCCGATTGCCACCATCGGACACACGGGTAGC | |
| RudS_KT_R60M | GGCTTGGGTGTCCCGATGGACCCGATTCGCCTG | |
| RudS_KT_D61M | GACCAGGCGAATCGGCATGCGCGGGACACCCAA | |
| RudS_KT_R64M | CCCGCGCGACCCGATTATGCTGGTCGGCAACCC | |
| RudS_KT_K110M | GGCCGCATGAAGGTGACATCTGCATGAAGATGTAG | |
| RudS_KT_S111M | CGTTCCAGGCCGCATGAAGGCATCTTCTGCATGAAGATGTAGC | |
| RudS_KT_S113M | CGTTCCAGGCCGCACATAGGTGACTTCTGCATGAAGATGTAGCC | |
| RudS_KT_E117M | CCTGATAAACCTTTACCCGCATCAGGCCGCATGAAGGTGAC | |
| RudS_KT_R118M | CCTGATAAACCTTTACCATTTCCAGGCCGCATGAAGGTGAC | |
| RudS_KT_K120M | TCCTGATAAACCATTACCCGTTCCAGGCCGC | |
| RudS_KT_Y122M | GGCCGTTGTCCTGCATAACCTTTACCCGTTCCAGGCCGC | |
| RudS_KT_H127M | TTATCAGGACAACGGCATGCCGGCCGTGCATGGTG | |
| RudS_KT_H131M | CCACCCGGCCGTGATGGGTGGCCGTGGCG | |
| RudS_KT_E152M | GCAGGCGGCCTTCCATTTCTACTGGCAGGTCCGGG | |
| RudS_KT_R155M | CCAGTAGAAGAAGAAGGCATGCTGCATGACCCTGTGCTG | |
| RudS_KT_H157M | GAAGAAGGCCGCCTGATGGACCCTGTGCTGCGC | |
| RudS_KT_N202M | CGATAGGCCTGGGGATTCATGGCCATCAGCAAATAC | |
| RudS_KT_N203M | GATAGGCCTGGGGCATGTTGGCCATCAGCAAATACTTGTA | |

| | |
|---|---|
| RudS_KT_Q205M | GATGGCCAACAATCCCATGGCCTATCGCACCCTC |
| RudS_KT_R240M | GCAGGCCCTGCGCATGTGCGCCAGCCGCG |
| RudS_KT_C241M | GCCGCGGCTGGCCATGCGGCGCAGGGC |
| RudS_KT_S243M | GTGCCGCGCATGGCGCAGCGGCGCA |
| RudS_KT_R244M | CGCCGCTGCGCCAGCATGGGCACCCACAGTAAC |
| RudS_KT_T246M | GCCAGCCGCGGCATGCACAGTAACGTGC |

**RESULTS**

**Supplementary table 4.** Groups of amino acids that are close to tRNA and might play a role in enzymatic catalysis and/or substrate binding.

| G1 | G2 | G3 | G4 | G5 | G6 |
|---|---|---|---|---|---|
| Arg24 | Arg60 | Lys110 | Glu152 | Asn202 | Arg240 |
| Tyr25 | Asp61 | Glu117 | Arg155 | Asn203 | Cys241 |
| Asn26 | Arg64 | Arg118 | His157 | Gln205 | Ser243 |
| Gly27 | | Lys120 | | | Arg244 |
| His29 | | Tyr122 | | | Thr246 |
| Lys30 | | His127 | | | |
| Asp33 | | His131 | | | |
| Arg36 | | | | | |
| Lys37 | | | | | |

**Detailed description of molecular dynamics simulations**

The systems underwent minimization, heating to 300 K, and equilibration. Constant temperature and volume production runs (NTV) were conducted at 300 K using Langevin dynamics thermostat. For the initial simulations of the holoenzyme and complex, during heating and the first equilibration run of 200 ns, Cartesian positional restraints of 1 kcal/mol/ $Å^2$ on backbone atoms of amino acids within ~7 Å around the iron-sulfur cluster were applied. These restraints were relaxed during the next 200 ns, and subsequent molecular dynamics simulations involved only distance restraints as specified. The same conditions were used for tRNA simulations.

The top HADOCK complex underwent a series of short molecular dynamics simulations, with 1 kcal/mol/ Å$^2$ restraints on protein backbone atoms and 5 kcal/mol restraints on the distance between the sulfur atom in the thiouridine moiety and the relevant iron atom. The distance restraint was incrementally reduced by one angstrom in each subsequent MD run until reaching a distance of 2 Å. The resultant complex then underwent 10 iterations of MD simulated annealing runs, each comprising 4 ns at temperatures ramping from 300 K to 400 K, 4 ns at 400 K, 4 ns cooling from 400 K to 250 K, 4 ns heating back to 350 K, 4 ns cooling back to 300 K, and finally 4 ns at 300 K, with restraints of 1 kcal/mol on protein backbone atoms and 5 kcal/mol/ Å$^2$ on the distance between the sulfur atom in the thiouridine moiety and the relevant iron atom. The final complex then underwent two different MD simulations, with the first one maintaining only 5 kcal/mol/ Å$^2$ restraint on the distance between the sulfur atom in the thiouridine moiety and the relevant iron atom, and the second one with an additional 4 distance restraints on specific Arg60, Glu51, Ser111, Ser113 atoms.

The best HDOCK model was used as a starting point in attempts to obtain a better model of the enzyme-substrate complex. Short molecular dynamics simulations with incrementally decreasing distance constraints for thiouridine sulfur and the relevant iron ion in the iron-sulfur cluster were conducted to bring the substrate and enzyme closer together. The obtained structures were then relaxed in 10 simulated annealing runs with previously described distance constraints and weak positional constraints on catalytic domain atoms within 7 angstroms around the iron-sulfur cluster. The resultant complex underwent initially restrained and restraint relaxation MD simulations as described in Methods and Materials, followed by a 640 ns simulation with the following 5 kcal/mol/ Å$^2$ distance constraints: Arg60 epsilon NH – Glu51 side chain carboxy O2; Glu51 side chain carboxy O1 – Ser113 side chain hydroxy group H; Ser113 side chain hydroxy group O – Ser111 side chain hydroxy group O, and thiouridine S – iron-sulfur cluster relevant iron ion. The last frame of this restrained simulation was used as an initial structure for two 260 ns simulations, where structure 1 retained the described above restraints, and structure 2 had only thiouridine S – iron-sulfur cluster restraint.

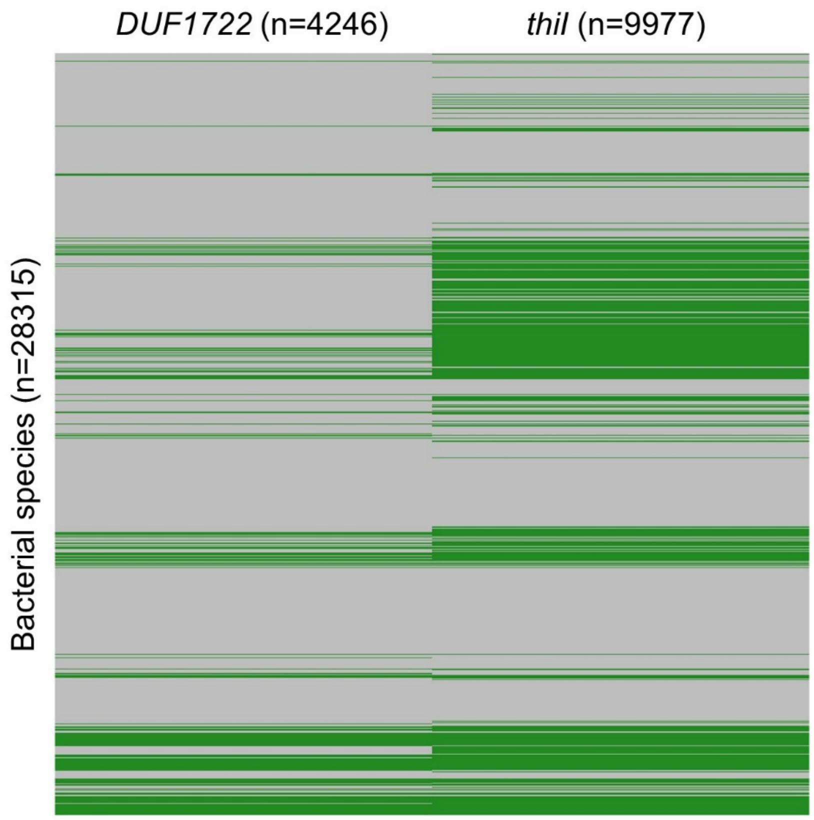

**Supplementary Figure S1.** Array representing phylogenomic distribution of genes among 28315 bacterial species (y axis). DUF1722 encoding species (n=4246, left panel) and ThiI encoding species (n=9977, right panel) are highlighted in green. The occurrence of both genes overlaps in 3993 species.

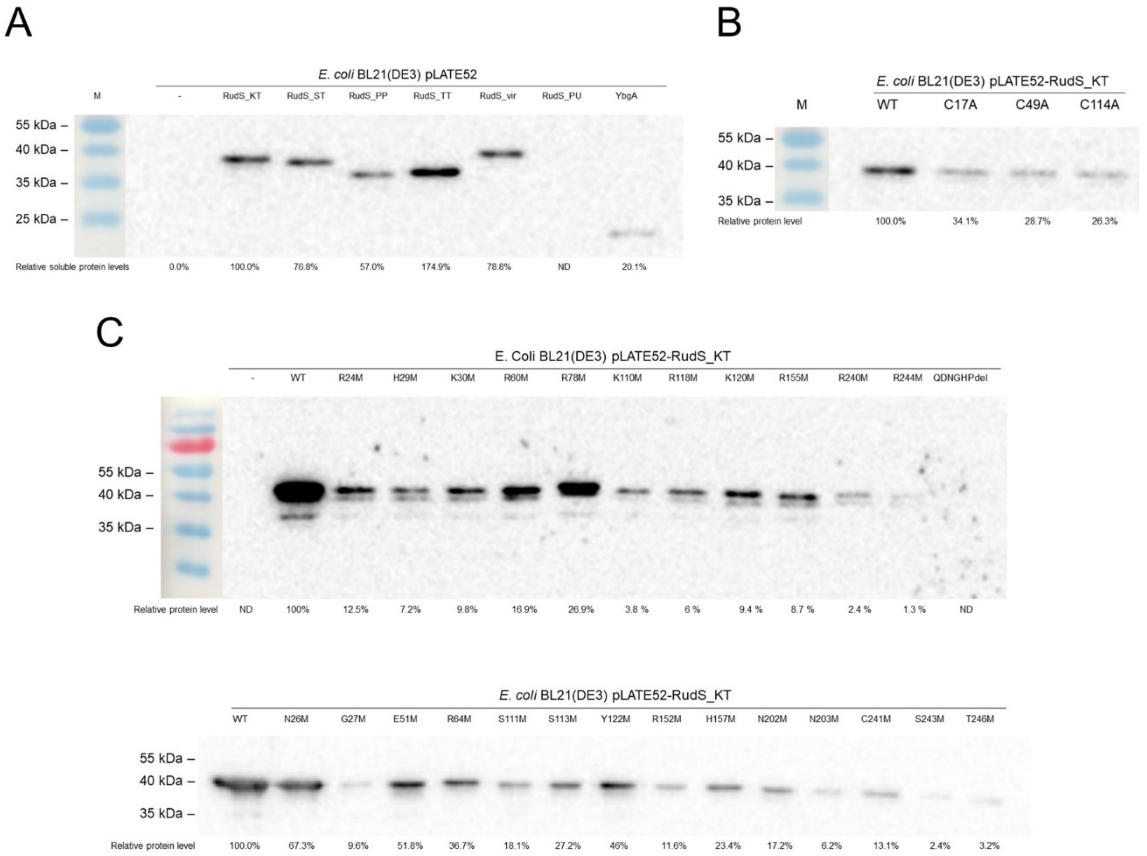

**Supplementary Figure S2.** Western blot analysis of soluble protein fractions in bacterial lysates (2 µg per lane). A: TudS-DUF1722 and DUF1722 (YbgA) domain containing proteins. B: RudS_KT mutants with substituted cysteines involved in iron-sulfur cluster formation. C: RudS_KT mutants having affected in vivo activity and predicted to participate in catalysis and/or substrate binding. Relative protein levels are normalized to wild type RudS_KT.

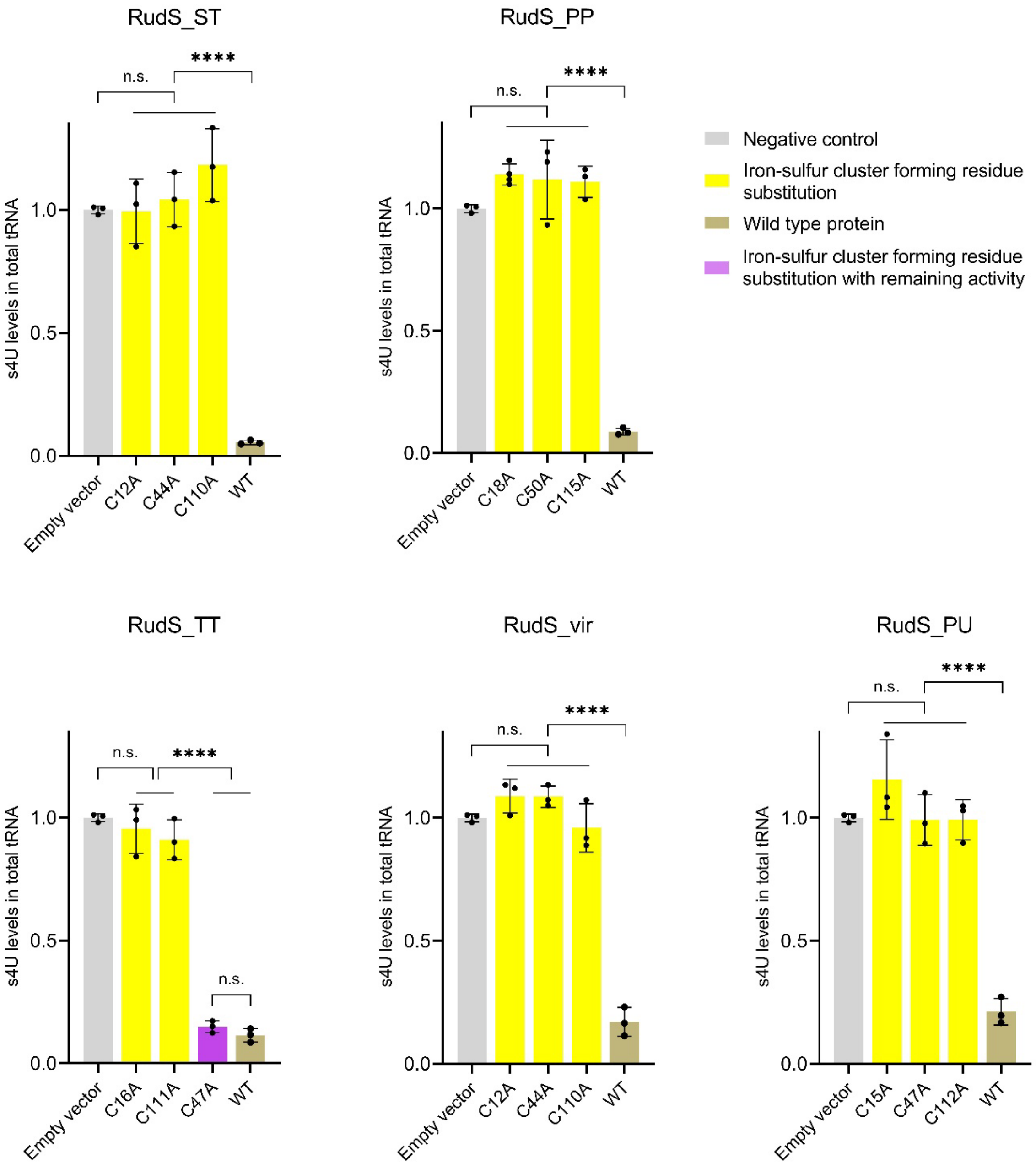

**Supplementary Figure S3.** s4U levels in *E. coli* total tRNA upon expression of RudS with conserved cysteine substitutions. ****: $p<0.0001$, n.s.: $p \geq 0.05$ compared to WT, one way ANOVA with Bonferroni's post hoc test.

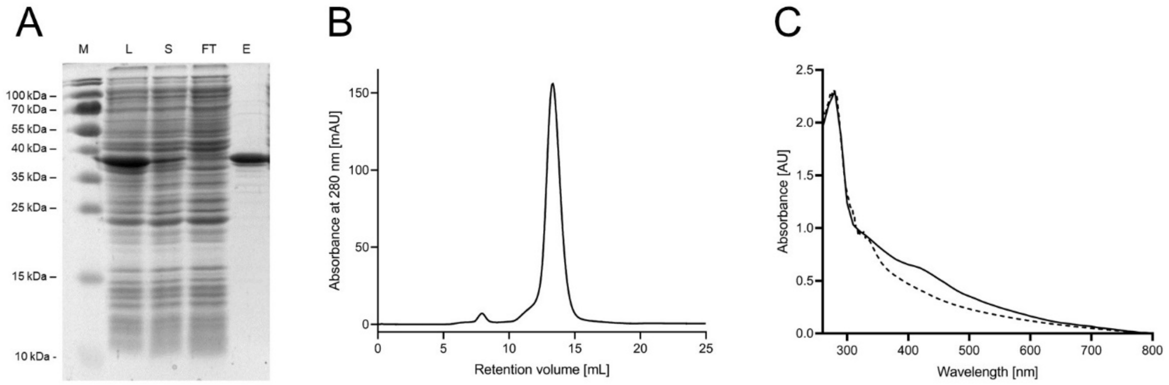

**Supplementary Figure S4.** A: Recombinant N-6xHis tagged RudS_KT purification using affinity chromatography (M – molecular marker, L – bacterial lysate, S – soluble fraction, FT- flow through, E – eluted protein; theoretical mass 39.21 kDa). B: Elution peak of purified RudS_KT after Superose 12 10/300 GL gel filtration chromatography (mass 37.1 ± 0.7 kDa). C: bold graph – UV-vis spectra of RudS_KT as purified (35 μM); dotted graph – RudS_KT after iron-sulfur cluster reduction with 10 mM sodium dithionite.

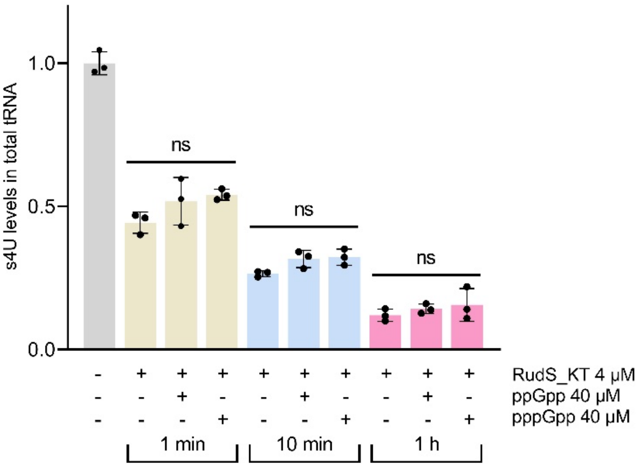

**Supplementary Figure S5.** Effect of guanosine tetraphosphate and pentaphosphate (ppGpp and pppGpp respectively) on RudS_KT in vitro activity. Three samples obtained within a single time point were compared to each other using one way ANOVA with Bonferroni's post hoc test (n.s.: p⩾0.05).

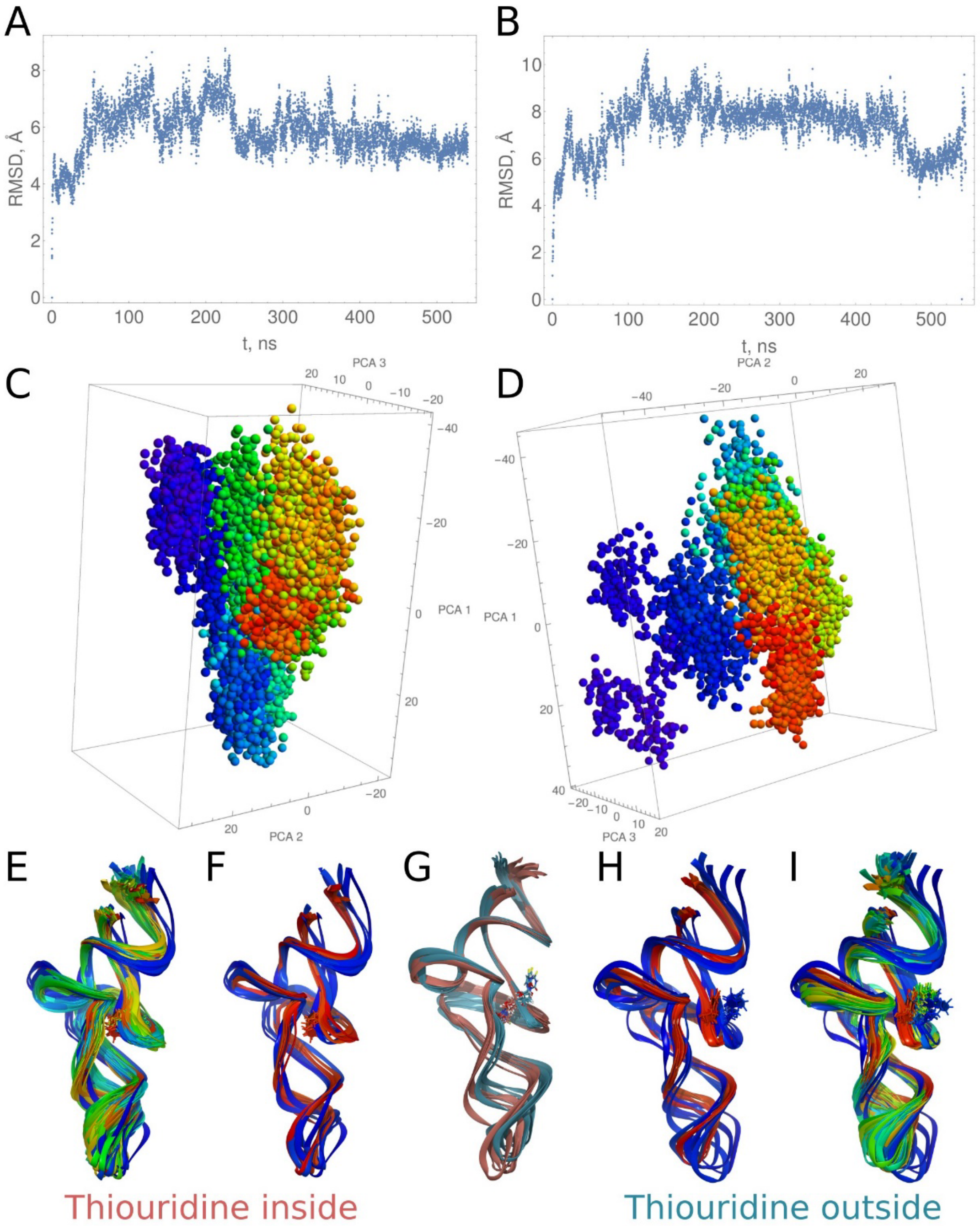

**Supplementary Figure S6.** Molecular dynamics simulation of phenylalanine tRNA from *E. coli* (PDBID:6y3g). A and B: RMSD plot of MD simulations of native and flipped out thiouridine harboring tRNAs. C and D: PCA analysis of MD simulations of native and flipped out thiouridine harboring tRNAE and I: overall motion of respective tRNAs from MD start (blue) to the end(red). F and H: overall motion of respective tRNAs from MD start (blue) to the end(red) with intermediate frames removed. G: superposition of final MD frames of native and flipped out thiouridine harboring tRNA.

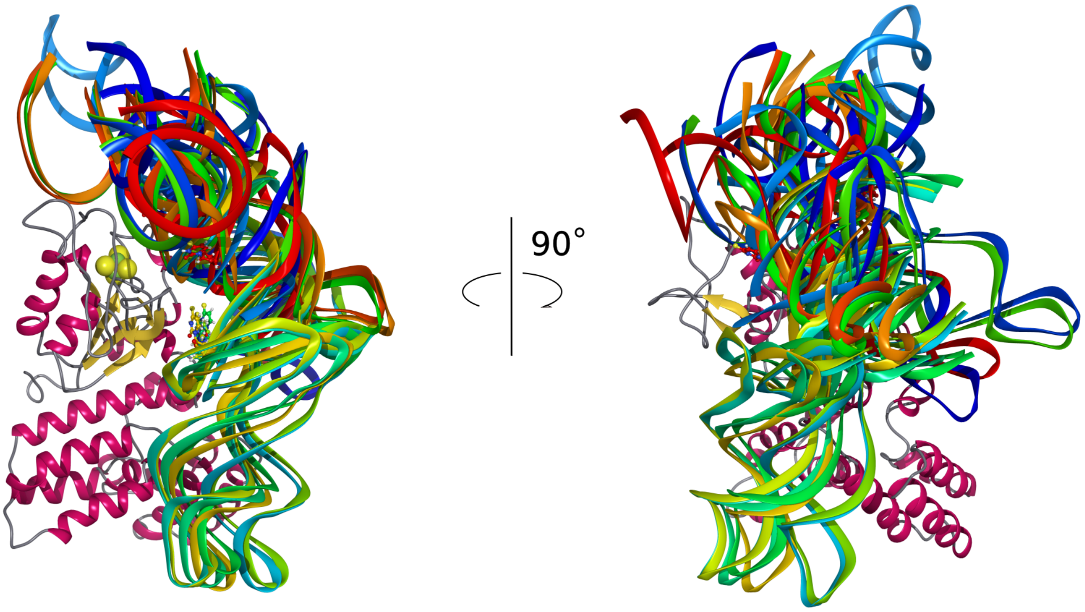

**Supplementary Figure S7.** Top 15 complexes produced by docking tRNA with thiouridine flipped out to the holoenzyme with HDOCK.

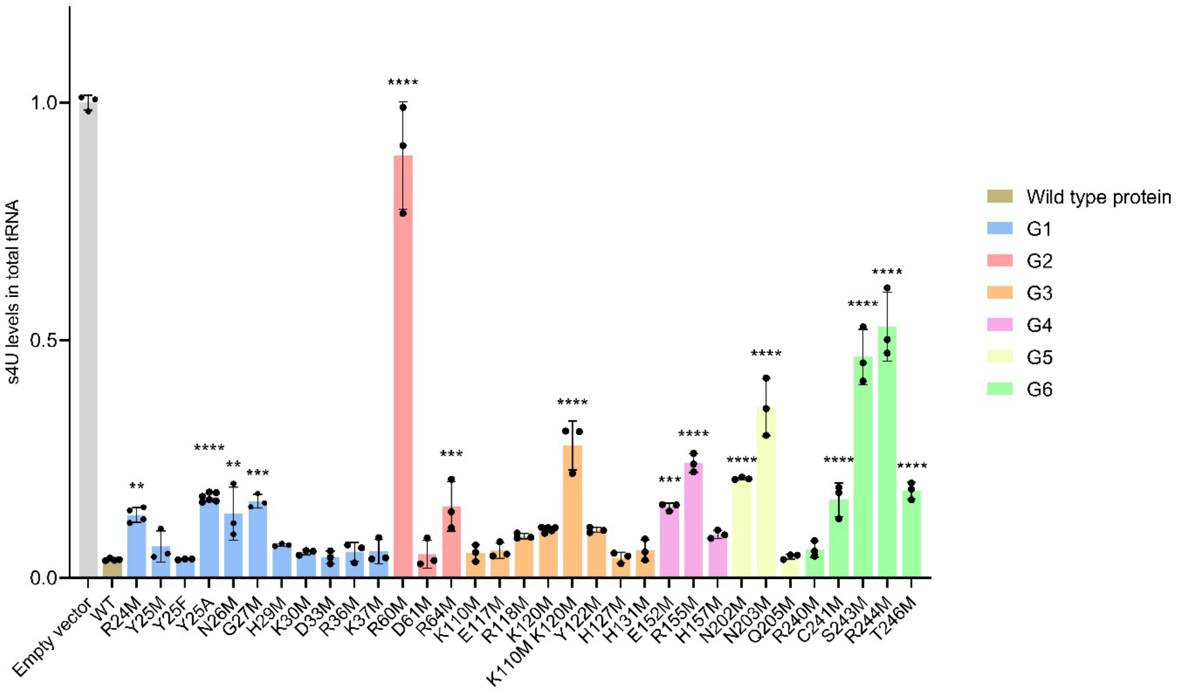

**Supplementary Figure S8.** s4U levels in *E. coli* total tRNA upon expression of RudS_KT with single amino acid substitutions. ****: $p<0.0001$, ***: $p<0.001$, **: $p<0.01$, *: $p<0.05$, n.s.: $p\geqslant 0.05$ compared to WT, one way ANOVA with Dunnet's post hoc test.

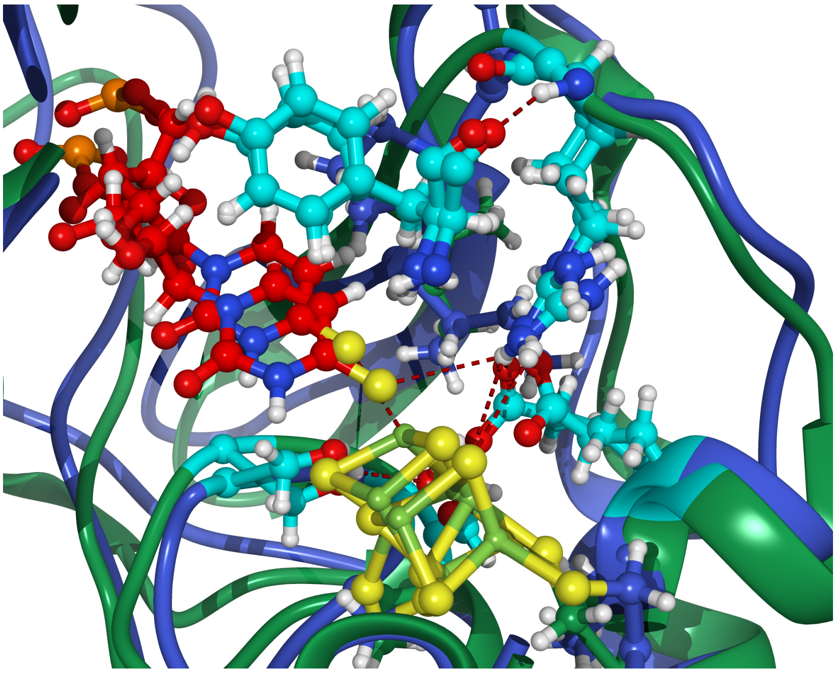

**Supplementary Figure S9.** The comparison of randomly selected frames from model 2 and model 1 molecular dynamics shows that orientations of catalytic amino acids relative to the thiouridine moiety are very similar in both models.

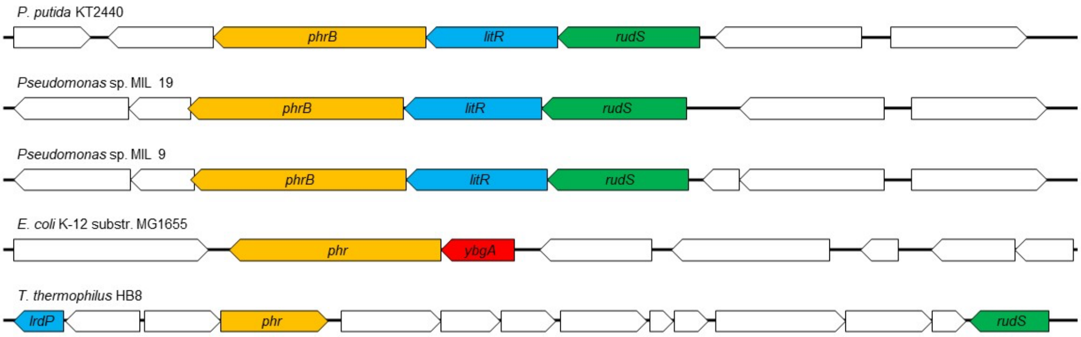

**Supplementary Figure S10.** Organization of RudS and YbgA encoding genome loci. Genes highlighted in green – RudS encoding gene; red – DUF1722 encoding gene; orange – DNA photolyase encoding genes; blue – light-inducible transcription regulator encoding genes.
